# Probabilistic Cognitive State Modeling (PCSM): Decoding Latent Spatiotemporal Dynamics to Reveal Serial-Parallel Processing, Cognitive Demand, and Serial Bottleneck in Task-Based fMRI

**DOI:** 10.1101/2025.10.31.685855

**Authors:** Drew E. Winters

**Author notes:** Corresponding author: Drew Winters.

## Abstract

Studying flexible, adaptive transitions between cognitive tasks and serial–parallel processing under changing task demands has been central to understanding human cognition. Advances in neuroimaging analysis have improved the ability to link cognition with brain function, motivating methods that characterize dynamic brain activity to quantify emergent cognitive properties during task-based fMRI.

Probabilistic Cognitive State Modeling (PCSM) combines Finite Impulse Response modeling of BOLD activity with a Gaussian Mixture Model–Hidden Markov Model to infer recurring multivariate patterns of task-evoked BOLD responses across spatially distributed regions over time (“brain states”). From the resulting posterior structure, PCSM deterministically derives interpretable processing metrics, including serial–parallel deviation, cognitive demand, and serial bottleneck.

Data-informed generative simulations evaluated PCSM across systematically varied noise levels and transition regimes. Results show that PCSM reliably recovers latent structure (∼98% state-alignment accuracy under known generative conditions) and produces stable parameter estimates across simulation regimes. Threshold analyses identify reliable boundaries between parallel, mixed, and serial processing modes and recover expected relationships among demand, and bottleneck.

Together, these results demonstrate that PCSM provides a principled framework for characterizing dynamic task-evoked processing architectures and estimating individual-level cognitive dynamics from task-based fMRI, supporting future investigation of cognitive processing constraints across tasks and populations.

## 1 Introduction

### 1.1 Background

Human cognition is supported by dynamic processing architectures that range from *parallel processing*, where multiple computations are handled simultaneously, to *serial processing*, where computations are conducted sequentially [1-3]. These computationally distinct architectures are not static states, rather adaptive responses to varying task demands, context complexity, and internal constraints such as cognitive capacity and cognitive load [4, 5]. These dynamic processing mode transitions are central to adaptive cognition, reflecting the system’s attempt to optimize limited control resources by balancing flexibility, efficiency, and precision under changing demands [4, 6].

Switching between cognitive tasks (i.e., distinct operations performed) and processing modes (i.e., the way cognitive operations are carried out) increases cognitive demand and taxes cognitive resources [7]. Cognitive resources are finite with individual-specific limits [8], constraining the ability to sustain or shift between operations and processing modes. Empirical work demonstrates switching between tasks and modes each impose a cost, and these costs are compounded when both occur simultaneously [9-11], suggesting that resource capacity itself is dynamic rather than fixed. Thus, task and mode transitions depend on cognitive control to flexibly allocate limited resources in support of adaptive cognition [4, 12].

This interplay between resource availability and cognitive transitions is thought to underly key dimensions of psychiatric dysfunction. Across transdiagnostic internalizing/externalizing spectra, this relationship is reflected in alterations during cognitive control, socio-affective processing, reward sensitivity, and processing flexibility [13-18].

### 1.2 Challenges with Current approaches

Despite the relevance of these transitions, traditional reliance on behavioral metrics imposes limitations that constrain our ability to measure them, particularly at the individual level. Cognitive and psychiatric phenomena are inherently heterogenous [19], yet most current tools for measuring cognitive dynamics depend on aggregate behavioral metrics such as response time and accuracy. These task outputs, while informative, are low-dimensional, sensitive to measurement noise [20], and constrained by repetitive response patterns [21], making them ill-suited for decoding individual-specific fluctuations in processing mode.

Experimental paradigms designed to elicit serial or parallel processing (e.g., dual-task, psychological refractory period [PRP], task switching with preview) rely on static trial categorizations that do not reflect individual-specific dynamic transitions over time. Structural features of these tasks such as manipulation of stimulus onset asynchrony [PRP: 22, SOA; dual-task: 23] or preview interval [task-switching with preview: 24] are assumed to induce a binary serial or parallel strategy in the subsequent trial. This approach, like behavioral metrics, constrains detection of within-trial fluctuations and individual-specific dynamics in processing architecture, reinforcing an oversimplified view of processing modes that does not reflect how processing strategies evolve dynamically over time or differ across individuals.

In addition to their static assumptions, such paradigms do not explicitly measure processing modes within core cognitive domains, limiting ecological validity and theoretical generalizability. Cognitive domains including cognitive control and working memory involve both parallel and serial processing [4], yet the existing paradigms are optimized to elicit processing interference in tightly controlled conditions incompatible with tests of cognitive function. This disconnect of processing mode study from cognitive functions restricts understanding how serial and parallel processing emerge and fluctuate within domains most relevant to learning, decision-making, and psychiatric risk.

Individual-specific brain dynamics, from which cognition emerges [25], can be more precisely examined by leveraging the temporal and spatial resolution of task-based functional magnetic resonance imaging (fMRI). Although task performance is often summarized by accuracy, the same observed behavior can arise from distinct neural computations [26]. The rich temporal and spatial properties of fMRI capture distributed activity patterns that, even during the same observed behavior, reflect distinct underlying transitions between BOLD-derived neural patterns [27]. However, prior fMRI research on serial and parallel processing [e.g., 28, 29] inherit the same limitations found in behavioral paradigms: they rely on trial-averaged signals across broad cortical regions, assume static mappings between task condition and processing mode, and use task designs that limit generalizability to core cognitive functions [30]. As a result, these approaches obscure within-trial neural dynamics and impede identifying individual-specific transitions in processing architecture.

### 1.3 Modeling Dynamically with Probabilistic Cognitive State Modeling (PCSM)

To address these limitations, Probabilistic Cognitive State Modeling (PCSM) is introduced as a data-driven framework that leverages fMRI to decode individual-specific latent processing states and their transitions. Here, “individual-specific” refers to subject-level estimation of time-resolved posterior trajectories and derived processing metrics, rather than to between-subject differences in cognitive ability. PCSM is not intended as a competing method for dynamic brain state discovery (e.g., dynamic functional connectivity or sliding-window approaches), but as a framework for deriving theory-constrained cognitive processing metrics from probabilistic state-space representations. PCSM extends principles originally applied to single-neuron recordings in rhesus monkeys [30], where dynamic latent brain states were inferred from trial-specific neural activity, to the domain of human hemodynamic measurement. Here, “brain states” refer to recurring time-resolved, trial-aligned multivariate patterns of task-evoked BOLD responses, rather than static summaries of activation or covariance structure across an entire time series.

PCSM integrates Finite Impulse Response (FIR) modeling with a Gaussian Mixture Model-Hidden Markov Model (GMM-HMM) to infer how brain states evolve over trials, brain regions, and time. FIR modeling enables trial-resolved estimates of BOLD responses, preserving temporal precision without assuming a fixed hemodynamic shape. The GMM-HMM then learns latent spatiotemporal state dynamics directly from these FIR-derived signals, capturing both state and transition probabilities.

This spatiotemporal modeling infers latent brain states representing recurring multivariate patterns of task-evoked BOLD responses, which covary with task performance [31], but themselves not interpreted as cognitive processes. Similarly, these latent states are not a representation of serial-parallel processing or any other cognitive metric; rather, the probabilistic structure of node-level responses across states and time learned from the BOLD in GMM-HMM are used to derive metrics in PCSM. By leveraging these latent dynamics of the brain, PCSM applies computational formulas to decode emergent and individual-specific properties in cognitive processing and demand. This allows researchers to track individual specific within-trial transitions from parallel to serial processing and cognitive load, quantify their stability temporally, and identify the contributions of distributed brain systems to processing transitions and cognitive resource fluctuations.

Consistent with contemporary understanding of BOLD and processing [32], cognitive processing is operationalized as temporally and spatially distributed changes in BOLD signal reflecting encoding, integration, and selection of task-relevant information. Processing is considered parallel when multiple brain regions show moderate levels of engagement at the same time, suggesting a distributed commitment across latent states that allows for concurrent handling of information streams. In contrast, processing is considered serial when engagement is more polarized, with regions strongly committed either to responding or not responding, reflecting a more selective, one-stream-at-a-time mode of operation. Within PCSM, these distinctions are captured by metrics at the node level (BOLD response probability by node: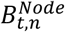) that, when aggregated, are used for inferring overall cognitive processing temporally (Serial-Parallel deviation: 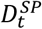). These metrics translate latent brain states into interpretable measures of how the brain allocates commitment over time.

### 1.4 Theoretical Background

Building on the logic of the psychological refractory period (PRP) framework, where structural or strategic bottlenecks constrain processing to occur in sequence [22, 24], PCSM extends this principal from behavior into human brain dynamics. Traditionally, serial and parallel processing were inferred indirectly through reaction time delays under varying stimulus onset asynchronies (SOAs), which are thought to induce serial or parallel processing for static classification. However, the core idea of PRP – that cognitive systems undergo structural or strategic constraints that prevent full parallel processing, especially under high control demands [24] – isn’t restricted to SOAs or static classifications. Therefore, this PRP principle can be generalized to dynamic bottlenecks observable in neural data, where demand and resource constraints shape the flow of cognitive processing over time.

In PCSM, the central processing bottleneck is extended to BOLD-derived neural dynamics. Serial processing emerges when a latent state reflects strong, polarized responses across regions, indicating functional commitment to a single process stream. Parallel-like processing emerges when latent states reflect moderate, distributed responses across regions, supporting simultaneous processing streams (Figure 1). Behavioral evidence shows that switching between tasks or processing modes imposes additional cognitive costs [9-11], consistent with PCSM’s quantification of dynamic demand and reallocation of cognitive resources.

**Figure 1.**
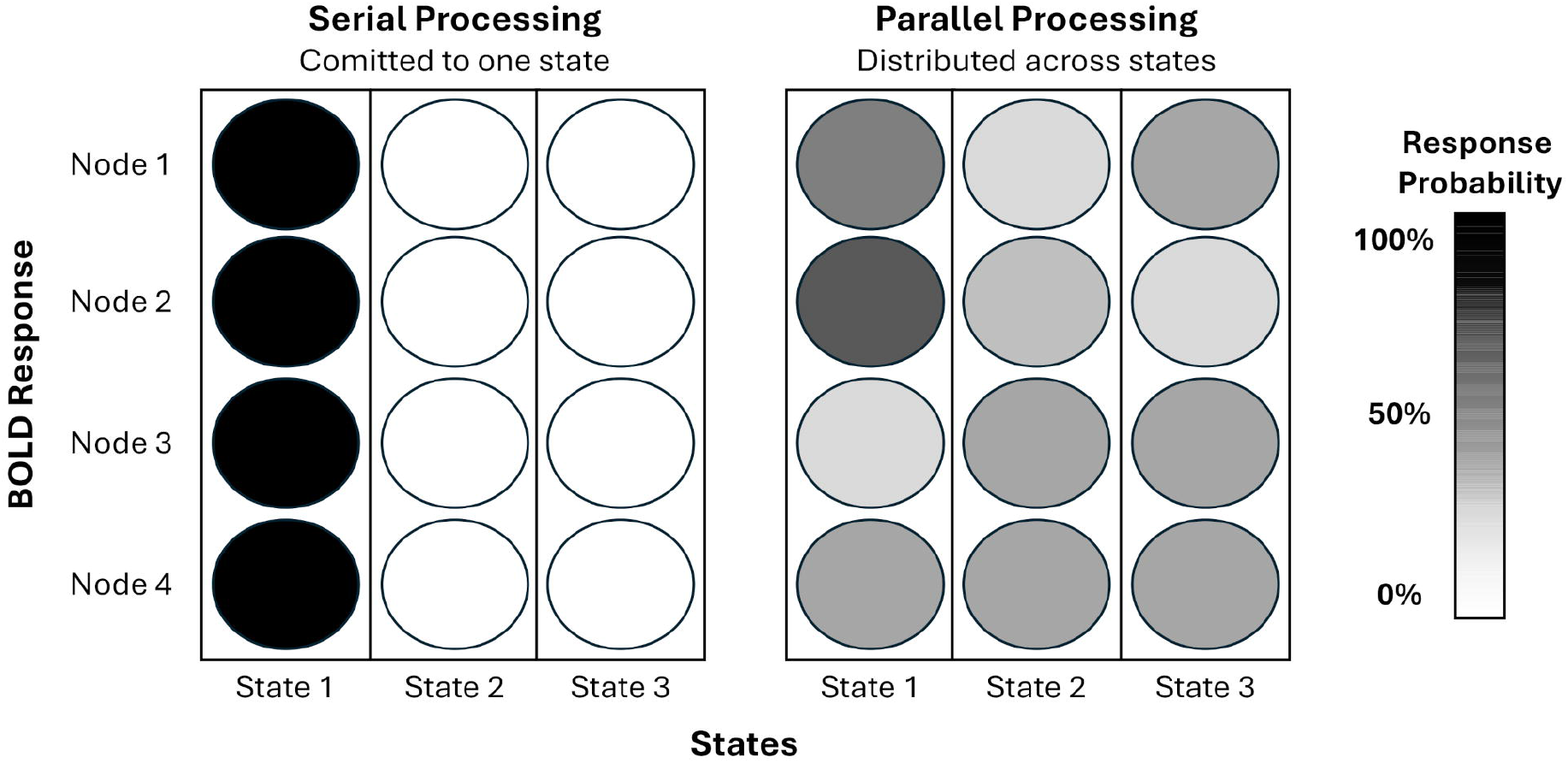

Traditional SOA approaches treated timing delays as indicators of serial persistence and exit difficulty. PCSM leverages dynamic fMRI data to model and quantify serial persistence and exit difficulty directly from BOLD signal, where the magnitude of individual-specific bottleneck effect is defined as a function of time spent in serial mode, cognitive demands, and the cost of switching. This reconceptualization enables serial and parallel processing to be inferred probabilistically and dynamically from fMRI data across a wide range of cognitive tasks. Thus, PCSM generalizes the PRP principle beyond SOA-based behavioral tasks to the spatiotemporal organization of human brain activity during any cognitive task.

### 1.5 From Theory to Unified Computational Model

PCSM is a unified computational framework with a modular composition that is designed to derive time-resolved processing metrics from task-based fMRI. It is not a method for discovering functional networks, estimating connectivity, or assigning cognitive meaning to latent states themselves. The unified modular framework proceeds in two stages (Figure 2).

**Figure 2.**
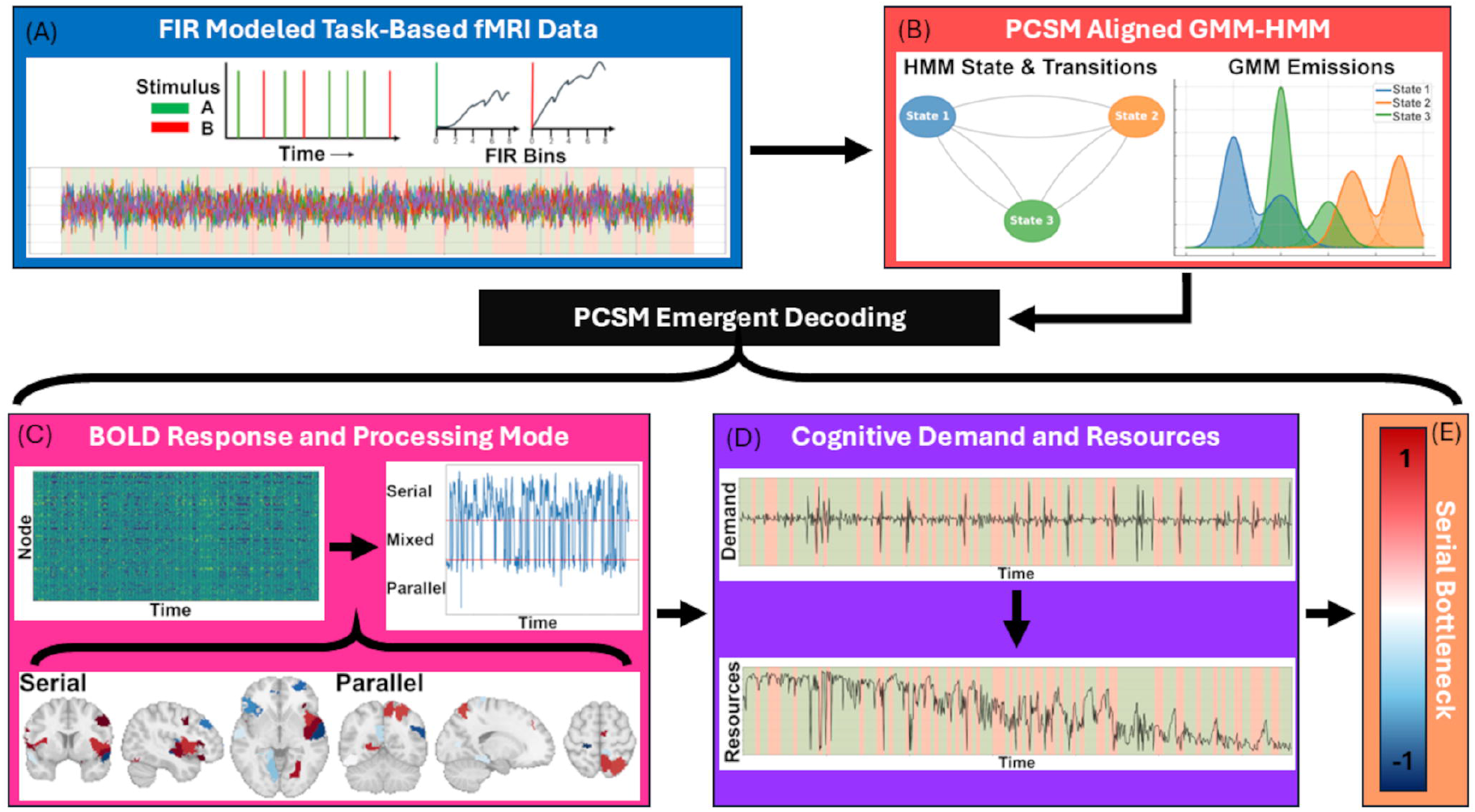

In the first stage, trial-aligned FIR modeling followed by a GMM-HMM yields posterior distributions over recurring multivariate patterns of task-evoked BOLD responses (“brain states”) and node-level response probabilities conditioned on state. Probabilistic mixture-based HMMs have been widely used in neuroimaging to model recurring multivariate neural configurations and their temporal evolution, including task-evoked activation patterns and dynamic neural states, precisely because they provide uncertainty-aware inference over heterogeneous response patterns while enforcing temporal coherence [e.g., 33, 34, 35]. In PCSM, the GMM-HMM is used to estimate the joint probabilistic structure of latent state occupancy and node-level response probabilities, constituting the fundamental computational object of PCSM.

In the second stage, cognitive processing metrics (e.g., serial–parallel deviation, demand/resource indices, bottleneck magnitude) are deterministically derived from the probabilistic posterior structure inferred by the GMM-HMM. Importantly, PCSM does not map BOLD signals directly to cognitive constructs; rather, cognitive features are formally defined functions of probabilistic response structure. Explicit mathematical derivations of each metric are provided in Section 2.1.4.2.

Under this formulation, each component of PCSM is computationally necessary: without FIR modeling, within-trial temporal specificity is lost; without probabilistic state-space inference, node-level response probabilities conditioned on brain-state configurations are ill-defined; and without joint probabilistic structure across nodes, constructs such as demand and bottleneck cannot be operationalized. PCSM therefore provides a principled and theory-constrained bridge between hemodynamic measurements and canonical models of cognitive processing limits.

While PCSM is motivated by questions central to computational psychiatry, the present work focuses on framework definition and methodological validation; empirical evaluation of psychiatric relevance is a key direction for future applications rather than a claim addressed in this study.

## 2 Methods

The preregistration for PCSM development, along with simulated data and its derivatives, can be found at the Open Science Framework (https://doi.org/10.17605/OSF.IO/DFJSB). All code used for simulation and recovery as well as the functions making up the PCSM package can be found in the projects GitHub repository (https://github.com/drewwint/pcsm).

Empirical human data were obtained from the open-access dataset OpenNeuro ds000030 (https://openfmri.org/dataset/ds000030/). All original data collection procedures were approved by the Institutional Review Board at the University of Texas at Austin, and informed consent was obtained from all participants [36]. The present secondary analysis complied with the ethical standards of the Declaration of Helsinki and did not require additional institutional approval. Analyses focused on the stop-signal task (trial-based event-related design). The stop signal task conducted in the fMRI scanner consisted of 259 subjects that completed 128 trials (75% go, 25% stop) where 184 volumes were acquired with TR = 2s, TE = 30ms, flip angle = 90o, voxel size = 3×3x4mm, and scan duration ≈ 6min 8s.

PCSM is optimized for trial-based task fMRI designs, where discrete events allow the model to capture rapid transitions between latent brain states. Block designs provide less temporal resolution for modeling these transitions. Thus, all analyses and validation were conducted on trial-based task fMRI data and simulations.

### 2.1 PCSM Pipeline

PCSM starts with Finite Impulse Response (FIR) modeling of each trials bold signal that is input into a GMM-HMM with PCSM defined alignment of states and mixtures. The aligned outputs are then used for diagnostics and emergent decoding metrics of node response, processing mode, cognitive demands, resource level, and serial bottleneck severity. These values can then be used to test any number of scientific hypotheses where cognitive processing, demand, and bottleneck are pertinent.

#### 2.1.1 Finite Impulse Response (FIR) BOLD Modeling

Nonparametric estimation of BOLD with the standard FIR [37] affords a fine-grained temporal resolution necessary for achieving the goals of PCSM. Other models including canonical or HRF models (e.g., gamma, double-gamma) impose assumptions on HRF shape that can mischaracterize effect estimates and introduce bias, particularly when individual variability in HRF timing or amplitude is important [38, 39]. Thus, the approach most suitable for the goals of PCSM is BOLD modeled with FIR.

FIR modeling estimates BOLD at discrete timepoints following a stimulus, yielding a minimally biased and high-resolution representation of task-evoked activity [39]. Typically, the BOLD response peaks around six seconds post-stimulus and undershoots before fully recovering to baseline [40]. The period from onset through peak (roughly 0-8 seconds) contains the greatest task-related signal sensitive to active processing; on the other hand, the undershoot has a stronger association with physiologic noise [41]. To prioritize signals associated with active task engagement while avoiding contamination from post-peak physiological noise and HRF undershoot, PCSM models the BOLD response using FIR delays from onset to 8 seconds (e.g., if TR = 2s; 5 FIR delays models 0s to 8s in 2s increments).

Let *T ϵ* ℕ: total number of BOLD timepoint volumes, *Rϵ* ℕ: total number of task trials, *D ϵ* ℕ: total number of FIR delays per trial (e.g., 5 delays 0-8s), *K* = *R× D ϵ* ℕ: total number of FIR regressors, *N ϵ* ℕ: total number of spatially distinct nodes, *Y ϵ* ℝ^*T×N*^: observed BOLD over time and space, *X ϵ* ℝ^*T×K*^: FIR design matrix, and 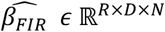: estimated FIR amplitudes. The FIR estimates are derived using the standard solution:

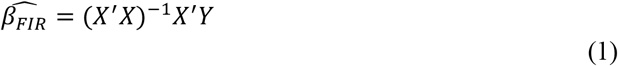

Where’ denotes transposition. This results in a trial- and delay-specific tensor of FIR responses with the shape *R× D × N*, which is reshaped into a matrix *Y*_*FIR*_∈ ℝ^*K×N*^, where each row corresponds to one indexed timepoint *t ϵ* &#x2124;_≥0_. With *y*_*t*_ = *Y*_*FIR*_[*t*,:] ∈ ℝ^*N*^, the the following is used as an input for GMM-HMM decoding:

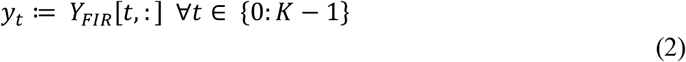

#### 2.1.2 Gaussian Mixture Model – Hidden Markov Model (GMM-HMM)

The GMM-HMM framework is leveraged by PCSM to robustly model multimodal spatial patterns and temporal dynamics of FIR-derived BOLD signals for individual-specific estimates. Here, the GMM-HMM is used as a statistical workhorse to obtain temporally coherent posterior distributions over brain states and node-level responses, which serve as inputs to downstream metric derivation rather than as endpoints of analysis. Distinct brain patterns reflecting brain states involve dynamic, spatially distributed, multimodal signals (e.g., respond and non-respond) that infer latent brain states, which both evolve and transition temporally [e.g., 42, 43-45]. These brain states are explicitly modeled by combining a Gaussian Mixture Model (GMM) with a Hidden Markov Model (HMM), leveraging their complementary strengths to redress limitations of each method alone.

The GMM captures the multimodal spatial patterns of BOLD responses at each timepoint, but does not account for the temporal dependencies inherent in fMRI data [46]. Conversely, the HMM models the temporal evolution and probabilistic transitions between latent states, but typically assumes unimodal emission distributions that may inadequately capture the complex, multimodal nature of fMRI signals within each state [46].

To redress these challenges, PCSM models the emission distributions of FIR-derived BOLD signals as Gaussian mixtures within each latent state of the HMM. Specifically, the emission probability of observing the multivariate BOLD vector *y*_*t*_ ∈ ℝ^*N*^ given latent state at time *t* is *C*_*t*_ = *c* for *c ϵ*{0,1,2} is modeled as a mixture of Gaussians *M* indexed *m ϵ* {0,1} :

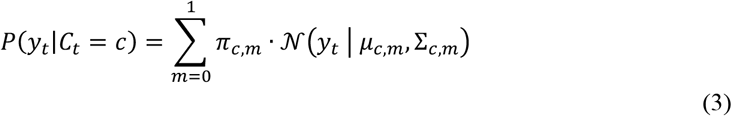

The mixtures are comprised of: *π*_*c,m*_= weight of the *m*^*th*^ Gaussian in state *c, μ*_*c,m*_= mean of the *m*^*th*^ Gaussian in state *c*, and Σ_*c,m*_= covariance matrices of the *m*^*th*^ Gaussian in state *c*. Model parameters, including the emission mixtures and state transition probabilities, are estimated iteratively using the *Baum-Welch algorithm* [47], an instance of the Expectation-Maximization (EM) framework tailored for Hidden Markov Models. The Baum-Welch algorithm alternates between two steps: computing the expected sufficient statistics of the latent states (E-step) and maximizing the expected complete-data likelihood by updating parameters (M-step). This process continues until convergence, resulting in robust estimates of emission distributions and transition dynamics within latent states and their temporal evolution.

For decoding, the Viterbi algorithm [48] is used to find the most likely sequence of latent states (the Viterbi path) that best explains the observed FIR-derived BOLD signals. This provides trial-level inference of brain states, enabling interpretable latent state trajectories across time.

Together, these algorithms [implemented via a fork of ‘hmmlearn’; 49] enable PCSM’s GMM-HMM to flexibly represent complex brain activation patterns as mixtures within temporally evolving brain states, balancing probabilistic state assignment with precise temporal decoding for individual-specific estimates.

##### 2.1.2.1 PCSM-Specific Adaptations

Modeling defaults for the number of Gaussian mixtures and latent states in PCSM are supported by theory and empirical literature that was further verified here (see section 3. Results). These defaults enhance interpretability and cross-study consistency.

###### First

PCSM defaults to two Gaussian mixtures (*M* = 2) per latent state in the GMM emission model, representing response and non-response patterns. Responding BOLD signals are characterized by both positive and negative values, reflecting increases and decreases in BOLD activity [50-52]. In contrast, non-responding regions exhibit BOLD signals centered around and near zero [53]. Prior neuroimaging studies have empirically justified modeling BOLD responses with two mixture components [46], consistent with electrical recordings in animals [30] and human BOLD time series [54, 55].

###### Second

PCSM defaults to three latent brain states (*C* = 3) representing cognitive operations during the fMRI paradigm of interest. This choice is informed by prior fMRI studies demonstrating 2-5 latent states during rest [56] and theoretical work supporting three latent states being specific to identifying processing modes during cognitive tasks [30]. Verification tests conducted here on human cognitive fMRI (see section 2.2.2 Human Data) confirm that three latent states efficiency model brain data during cognitive tasks without unnecessary noise and complexity. Adding states beyond three yielded minimal gains in model likelihood while reducing interpretability, consistent with prior HMM-based cognitive-state research [35]. Therefore, three states are a feasible and verified assumption meeting the aims of PCSM that is applicable across a wide variety of tasks.

###### Third

PCSM incorporates three modeling adaptations to improve accuracy and stability. (1) *Incorporating temporal context*: consistent with Shappell, Caffo [56] and Fincham, Lee and Anderson [35], PCSM integrates FIR delay bins to inform the model for the duration spent in each trial phase. This preserves fine-grained temporal resolution. (2) *Latent state and mixture alignment across all individuals*: Due to stochasticity in GMM-HMM estimation can arbitrarily label or permute latent states and mixtures, PCSM aligns each subject to the global template by sorting states according to a salience core that combines (a) the state’s average BOLD intensity and (b) the state’s probability of being in the responding mixture (by either fixed mixture weights or subject-adjusted effective weights). The subject values are permuted across start probabilities, transitions, mixture weights, means and covariances to match the global state. This serves to preserve likelihood while ensuring consistent state labeling and interpretability across subjects. (3) *Fixing covariance matrices for numerical stability*: numerical instability arises from highly variable and extreme transition probabilities. PCSM constrains individual model covariance matrices to those estimated from a global concatenated dataset. This reduces model complexity while allowing other parameters (i.e., mixture means, mixture weights, and transition probabilities) to adapt at the individual level. This approach ensures stable estimation and meaningful subject-level decoding of dynamic brain states.

Together, these adaptations ensure stable estimation and meaningful comparability of individual-specific brain dynamics across participants. They also enhance the robustness and interpretability of PCSM’s individual-specific decoding of latent brain states.

#### 2.1.3 Values Extracted from GMM-HMM

From the GMM-HMM individual-level model fit, PCSM extracts the quantities at each timepoint *t* to quantify cognitive processing states. First, p*osterior state probabilities* are extracted: 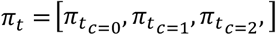, where *π*_*t*_,_*c*_ = *P*(*C*_*t*_ = *c*|*y*_0:*t*_) indicates posterior probability of latent state *c* and time *t*. Second, *mixture weights* by state are defined under the assumption that nodes *N*_*i*_ are conditionally independent given *C*_*t*_. The probability of responding (*m* = 1) given state *c* is extracted from the mixture weights matrix *A ϵ* ℝ^*C×M*^ = {*a*_*c,m*_} where each element is *a*_*c,m*_ = *P*(*M* = *m*| *C* = *c*). This is defined as *a*_*c*_ := *a*_*c,m*=1_ = *P*(*m* = 1| *C* = *c*), which quantifies the probability that a node *N*_*i*_ is responding when the system is in latent state *c*. These extracted values then are used by PCSM to compute a set of diagnostic and decoding metrics detailed below.

#### 2.1.4 Probability Metrics for Diagnostics and Emergent Decoding

To make probabilistic inferences on processing state, PCSM uses a set of cognitive metrics to derive interpretable and comparable values of emergent cognitive properties from GMM-HMM posterior state probabilities. Diagnostic metrics, ρ_*t*_, *P*_*t*_, and 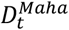, inspect temporal consistency, probability of response, and deviation from expected (respectively) for probing the temporal reliability to consider for processing state decoding. Processing mode metrics, 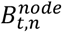 and 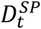, are used to temporally quantitate node level response probabilities that are used then to decode whole-brain and individual specific processing state (respectively). Cognitve demand 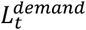, resources 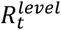, and the overall serial bottleneck *S*^*bottleneck*^ quantitate the individual-specific fluctuations in cognitive challenges and the impact on resources available that inform the overall degree of a serial bottleneck (respectively).

##### 2.1.4.1 Diagnostics Metrics

PCSM uses three diagnostic metrics to evaluate the quality and stability of decoded cognitive processing states, ensuring robust and interpretable inference: the brain response probability *P*_*t*_, temporal reliability ρ_*t*_, and model–data discrepancy 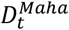. Together, these metrics characterize how faithfully the model captures the observed BOLD dynamics and identify timepoints that may be unreliable for inference.

###### Brain response probability *P*_*t*_

The instantaneous probability that the brain is actively responding to task stimuli. Where mixture component *m* = 1 indicates the response mixture, *a*_*c,m*=1_ = *P*(*M* = 1| *C* = *c*) is the state-specific mixture weight and *P*_*t*_ *ϵ*{0,1}, *P*_*t*_ is:

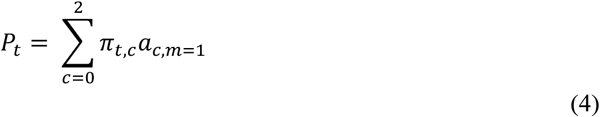

Extremely low values indicate where task-related BOLD activity is minimal, suggesting those periods may not reflect meaningful task engagement.

###### Temporal reliability metric *ρ*_*t*_

Stability of latent BOLD engagement over time ρ_*t*_*ϵ* {0,1} is defined as one minus the total variation distance between consecutive posteriors:

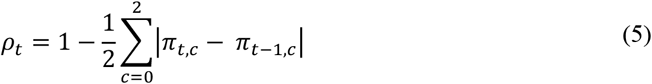

To improve signal to noise for temporal stability, smoothing via an adaptive exponential moving average is applied (optional) that does not change the definition. Here high values indicate reliable stability whereas low values indicate suggest period of unstable or less temporal reliability.

###### Model-data discrepancy 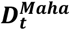

Mahalanobis distance metric 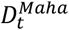 quantifies the difference between observed and model predicted BOLD. Let *y*_*t*_ *ϵ* ℝ^*N*^ be the observed BOLD at time *t* (across N nodes) and let *μ*_*c*_*ϵ* ℝ^*N*^ be the mixture-weighted mean for state c, then 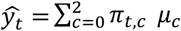. Where Σ*ϵ ℝ*^*NxN*^the spatial covariance across nodes, and Σ^−1^ is computed with a numerically stable inverse:

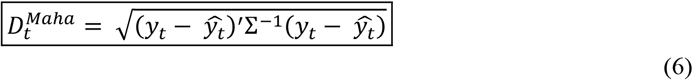

Higher 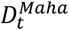 values indicate greater deviation from expected brain states, implying greater uncertainty in state decoding. Conversely, lower values suggest that observed BOLD activity closely matches predicted states, reflecting more reliable decoding.

###### Interpretation and Quality Control

Together, ρ_*t*_, *P*_*t*_, and 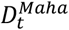 provide a comprehensive, multidimensional assessment of decoding reliability, BOLD response, and model fit, enabling identification of data segments that may compromise interpretability or require exclusion. Specifically, 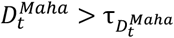 is used to identify timepoints to consider for further investigation with 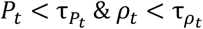 flagging frames for removal.

If a trial has > 50% of FIR bins flagged (i.e. 3 or more), then that trial is strongly recommended for removal. If < 50% of FIR bins are flagged (i.e., 2 or less), this suggests a strong transition that reaches stability within trial and recommended to keep that trial in. These diagnostic procedures ensure that state decoding reflects stable neural engagement rather than transient fluctuations or poor model fit.

##### 2.1.4.2 Emergent Decoding Metrics

###### Node response 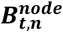

BOLD response probability by node 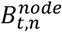 quantitates the posterior probability that a node *n* is responding at time *t*. Where *π*_*t,c*_ = ∑_*m*_ γ_*t,c,m*_ is the posterior probability of latent state *C* = *c* given the observations *y*_0:t_ and γ_*t,c,m*=1_(*n*) = *P*(*M*_*t*_ = 1|*y*_*t,n*_, *C*_*t*_ = *c*) is the mixture responsibility at node *n* for the responding mixture (noted as *m=1*) within state *c*:

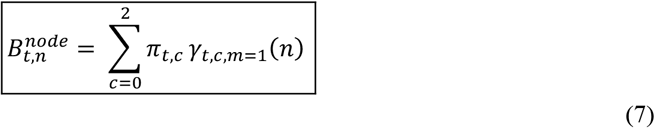

For Gaussian emissions with state-specific mixture weights *w*_*c,m*_, node-wise means *μ*_*c,m,n*_and variances *σ*^2^ :

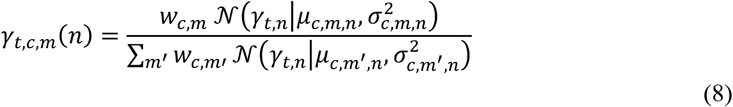

Thus 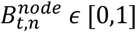 is a state-marginalized posterior of node-level responding, reflecting the likelihood that node *n* contributes to the responding mixture at time *t*.

###### Serial/parallel processing 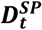

Whole brain processing mode is decoded from 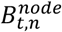 with the Serial-Parallel deviation index 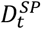, which quantitates the normalized deviation between the number of responding nodes (*K*_t_) and half the number of available nodes 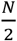 . Where *K*_t_ is a random variable representing the number of nodes at time *t, k ϵ K* is the realization of *K*, and *p*_*t*_(*k*) is the Poisson-binomial PMF:

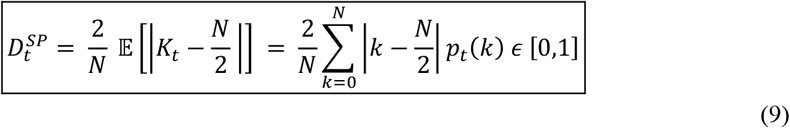

Where *p*_*t*_(*k*) is the Poisson-binomial PMF, let 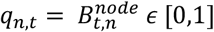 denote node-wise posterior probability of being in the responding mixture (denoted m=1) at time *t*:

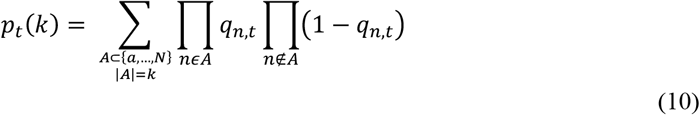

Higher values indicate serial processing (responding is concentrated on fewer nodes); lower values indicate parallel processing (responding is disbursed across many nodes). The factor 2/N normalizes the index across different N values. This is then later classified (parallel/mixed/serial) using data-driven thresholds (e.g., KDE valley minima) for reliable cross study comparisons.

###### Cognitive demand 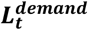

The signed moment-to-moment flow between processing strain and recovery 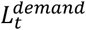 quantitates instantaneous cognitive load demand. It captures dynamic changes in processing difficulty arising from fluctuations in processing mode, latent-state uncertainty, and temporal transitions between brain states. Let 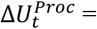 the normalized processing mode potential (derived from 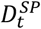); Δ*H*_*t*_ = the change in normalized posterior entropy of the latent-state distribution; *ΔJ*_*t*_ = the centered Jensen-Shannon divergence between successive posteriors (i.e., *y*_*t*_ − 1, *y*_*t*_); *S*_*t*_ = the median-centered synergy term (upweights steps with concurrent processing mode and cognitive state changes); and ω_s*ym*_ = the mean absolute magnitude of the unweighted demand, then instantaneous cognitive demand is:

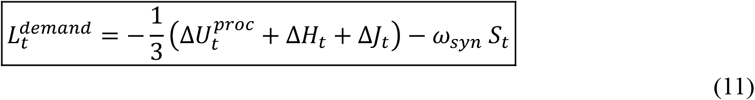

This metric is then scaled and normalized (median/MAD) with no intrinsic bound for use to derive the resource and bottleneck metrics as to not distort downstream calculations. To improve comparability of this singular metric, a hyperbolic tangent transformation (tanh) is applied 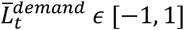. Here negative values indicate increased demand or difficulty, and positive values indicate recovery or stabilization.

###### Resource level 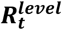

The cumulative availability of control resources follows a discrete-time leaky integrator 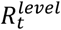 quantitates how resource levels accumulate and dissipate over time with individual-specific data-driven memory limits [e.g., 57, 58]. An exponential decay is applied on past contributions so that older demand values exert progressively less influence on the current resource state. Where λ is the leaky parameter determined from the individual-specific data-driven half-life (*Dwell*_*med*_ = median dwell time in processing modes) let *h* = max (3, 2*Dwell*_*med*_) then 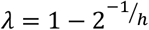; and where α is a data-driven resource scaling factor matching unit variance of 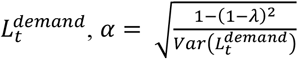 across subjects:

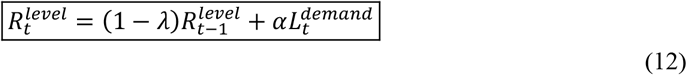

That is scaled to a bounded interval using robust normalization and hyperbolic tangent transformation (tanh) 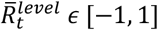 where higher values (*≈* 1) indicate more resource availability and lower values (*≈* −1) indicate a depletion of cognitive resources.

###### 2.1.4.2.3 Serial bottleneck

The serial bottleneck, *S*^*bottleneck*^, is a scalar value quantifying the persistence of serial processing and the effort required to disengage. Where *S*_s_ is the set of contiguous timepoints in the s^*th*^ serial episode (*S*_*tot*_ total episodes), *t*_*e*_ is the final timepoint of each serial episode with available post-exit data (*S*_*exit*_total), and *PEΔ* is a short post-exit interval (e.g., 3 TRs) with τ indexing timepoints in within that interval, this metric is derived by:

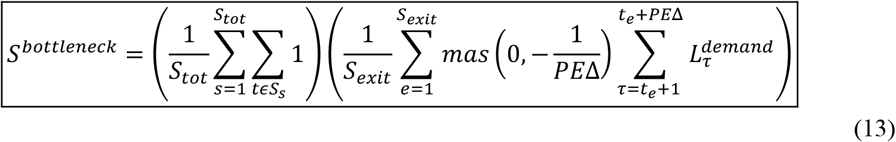

Thus, *S*^*bottleneck*^ integrates both persistence and exit costs to index the overall degree of serial constraint in cognitive processing. This metric is then robust scaled and hyperbolic tangent transformation 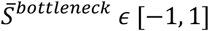. Here, higher values (*≈* 1) indicate longer serial persistence that is more difficult to switch from, consistent with a serial bottleneck, whereas lower values (*≈* −1) indicate a brief and easy transition serial bottleneck.

### 2.2 Simulation Design

To test identifiability and recovery of GMM-HMM parameters that form the computational basis for stable estimation of cognitive processing modes and PCSM metric derivation, a simulation with independently specified ground truth was designed for testing model recovery. Simulations were data-informed to ensure realistic signal, noise, and multivariate response structure. Empirical human data were used to guide the selection of latent state and mixture dimensionality and to calibrate scale and variability of simulated BOLD responses; however, simulated state sequences, emission parameters, and ground truth were defined by the generative model and not derived from empirical PCSM estimates or downstream processing metrics.

#### Task and Behavioral Simulation

The simulation models a two stimuli paradigm consistent with the stop-signal task (SST) incorporating key experimental and BOLD signal features. A sequence of go and stop trials is created with adjustable proportions, randomly spaced with inter-stimulus intervals (ISI) uniformly sampled within a specified range and normalized to fit the total simulated duration.

Following the SST consensus guidelines [59], the event-related SST simulation had a minority of stop trials (60% go and 40% stop), and there were sufficient number of trials for at least 50 stop trials (132 trials x 40% stop = 54 stop trials). Both reaction times and SSDs were sampled from a uniform distribution within realistic ranges. Outcomes (successful or unsuccessful stops) are determined by comparing stop success timing with go reaction time, generating a detailed trial-level behavioral data frame.

Ground truth cognitive state sequences, conditioned on trial type, are generated per trial with transition stability consisting of low, medium or high to simulate varying dynamics of cognitive state switching.

#### BOLD Signal Simulation

Given the simulated behavioral data, BOLD activity is simulated for 15,000 subjects across 200 spatial nodes for discrete FIR delay bins (0 to 8 seconds in 2-second increments for a total of 5 bins). This produced a three-dimensional tensor [trials × delays × nodes]. These nodes BOLD signals were weighed by state. Each latent cognitive state *c* ∈ {0,1,2} is associated with distinct node-specific activation patterns randomly initialized within amplitude ranges reflecting high positive, low positive, and mixed positive/negative activations, respectively. Gaussian noise was added to each node’s activity per trial and delay, with noise levels varying across four levels of noise (low, medium, high, very high) to simulate realistic signal-to-noise conditions. Simulated FIR-BOLD estimates are z-scored are returned along with the simulated SST behavioral data frame and the ground-truth latent states that are used for parameter recovery.

#### 2.2.1 GMM-HMM on Simulated data

The GMM-HMM was applied to the simulated dataset to evaluate the stability, recovery, and alignment of latent brain states under known ground-truth conditions. This step was critical for verifying that the model, priors, and emissions used in PCSM yield stable and identifiable latent state inference under known generative conditions.

Prior to recovery analysis, model parameterization was verified using realistic priors for state and mixture configurations and transitions probabilities (see section 2.2.2 Human Data). The same parameterization scheme was then applied to the simulated FIR-BOLD data to assess: parameter recovery accuracy for emission means, covariances, and mixture weights; state alignment stability across subjects and noise conditions; cross condition consistency in decoding of latent cognitive state transitions. This serves as the quantitative basis for assessing PCSM’s latent state recovery, ensuring its validity for downstream decoding of emergent decoding metrics.

#### 2.2.2 Human Data-Informed Model Selection

##### Cleaning and BOLD modeling

Determining the number of mixtures and latent states was jointly conducted with the The preregistration for PCSM development, along with simulated data and its derivatives, can be found at the Open Science Framework (https://doi.org/10.17605/OSF.IO/DFJSB). All code used for simulation and recovery as well as the functions making up the PCSM package can be found in the projects GitHub repository (https://github.com/drewwint/pcsm).

For all 259 participants in this dataset, raw data was cleaned using the specifications provided in this dataset and first level BOLD was modeled with FIR spanning bins up to 8 seconds (TR=2, 5 bins = 0-8s) after each stimulus and parcellated using the Schafer 200 atlas [60].

##### Evaluation Approach

Whole brain data was placed into the GMM-HMM model and evaluated with a two-pronged approach: (1) subject-level five-fold cross validation across 1-3 GMM mixtures within 2-5 HMM latent states and (2) whole sample five-fold cross validation of selected GMM-HMM parameters from subject level test. This two-pronged approach provides complementary evidence for selecting the most parsimonious generative model that generalizes across subjects in terms of likelihood and stability. Throughout, references to generalizability refer to the applicability of the PCSM framework and its computational definitions across tasks and datasets, rather than to predictive generalization of inferred metrics to held-out empirical data.

##### Model Selection Procedure

5-fold cross validation was conducted in scikit-learn [61] across a set of *M* = 1 − 3 and *C* = 2 − 5 latent states. This was consistent with work suggesting that 2-5 brain states generally [56] with three [30] and five [62] states expected to capture parallel-serial processing. In each fold, 80% of the subjects were used for training and 20% were held-out for testing. For each potential latent state *C*, the held-out log-likelihood per observation (*LL*_*c*_) and its standard error (*SE*_*LL*_) were computed across folds. Each model was compared to the next simpler model to assess information gain *ΔLL*_*C*→ *C*+1_ = *LL*_*C*+1_ − *LL*_*C*_.

##### Latent State Selection Criteria

Number of latent states were selected by (A) information gain criteria: the most parsimonious model with the most substantial gain over the prior model model that didn’t inflate error (ΔLL > 2 and ΔLL > 2 × *SE*_*LL*_ ; and for state changes, Δ*LL*_*C*_ > 0·10 *×* Δ*LL*_*C*−1_), and (B) consistency criteria: with the higher-order model outperforming the prior model in at least 70% of folds. Where applicable the next highest-order model had to improve information gain of at least 10% higher than the prior model.

##### Mixture Selection Criteria

For mixture selection, the same hierarchical approach was used. For each *C*, the number of Gaussian Mixtures *M* was tested in ascending order, comparing consecutive mixture configurations based on the held-out log-likelihood *LL*_*c*_(*M*). A higher mixture was retained only if its gain over the the previous mixture was practically meaningful indicated by *ΔLL*_*M*_ ≥ max (5, 2 *× SE*_*LL*_) per observation, yielding a single parsimonious solution for *C* and *M* for simulation parameters. This procedure evaluates out-of-sample likelihood to ensure stable and non-overfit generative model selection.

##### Model Selection

Results identified a three-state model with two gaussian mixtures met these criteria with real data (Table 1). This model configuration was used as the default parameterization for all subsequent simulations and ground-truth recovery analyses, without further tuning to empirical metric. While a three-state solution is used as a parsimonious default, PCSM does not depend on a fixed number of latent states, and higher-order solutions typically add rarely visited states without improving downstream metric estimation [35].

### 2.3 Parameter Recovery

#### 2.3.1 Cross-Validation of Global Model for Individual Empirical Priors

A two-pronged approach was used to evaluate the global model as a source of empirically informed priors. First, log-likelihood stability across folds was evaluated to determine robust explanation of unseen data. Criteria for log-likelihood stability was determined by all fold values remaining within 2SDs of the mean. Second, similarity between fold-specific and expected global model spatial patterns was evaluated for recovery of interpretable and generalizable spatial patterns from the global model, which was quantified by the Dice coefficient [63]. Dice values (0 = no consistency to 1= perfect consistency) quantified the alignment of fold-specific and global model patterns. This two-pronged approach provides a comprehensive evaluation of robustness, generalizability, and interpretability of PCSM.

#### 2.3.2 Latent State and Transition Recovery

To evaluate the accuracy of latent state and transition recovery, the model-inferred sequences were compared against ground-truth sequences form the simulation. For latent states, the correspondence between true and recovered states at each time point. For transitions, a binary variable was defined indicating whether the system remained in the same state or switched from the previous timepoint, which was compared to ground truth transitions. Recovery evaluations were conducted across noise levels and transition probabilities for overall expectations on model performance. Subsequent sensitivity analyses evaluations were examined by transition probability (high, medium, low) and noise level (low, medium, high, very high) to further evaluate under what circumstances that model recovery changes.

Recovery was evaluated using functions in scikit-learn [64]. Specifically, the classification_report function was used for precision, recall, F1-scores, and overall accuracy; and the confusion_matrix function was used to build a direct visualization of agreement between ground truth and model-inferred states. This approach ensured a transparent and standardized quantification of model performance across both levels of the latent process: discrete state recovery and dynamic transitions.

### 2.4 Determining Thresholds and Scaling Factors for Metrics

#### 2.4.1 Thresholds

Data-driven thresholds were estimated from the simulated distributions, which span a wide range of realistic transition patterns and noise levels. When a metric’s empirical density is multi-modal, Gaussian mixtures were fit for extracting density intersections between adjacent components as decision boundaries as follows. Where mixture with weights (*w*_1_, *w*_2_), means (*m*_1_, *m*_2_), and SDs (s_1_, s_2_), the intersections τ solves:

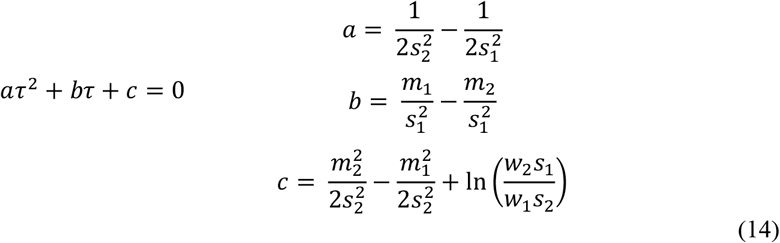

Mixture quality involved a mixture density > 20% and a separation ≥ 1, where <1 = weak, ∼1 = moderate, and >1.5 = strong) with a Cohen’s D like metric derived by:

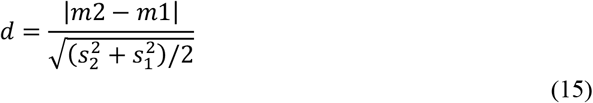

If a mixture quality is insufficient or the split is implausible for the observed shape, boundary-corrected KDE valley selection was applied. This involved smoothing a high-resolution histogram with a Gaussian kernel under reflection at the boundaries, and defining thresholds as local minima within pre-specified windows around expected density dips. Threshold stability was further assessed through sensitivity analyses varying histogram resolution and kernel bandwidth with drift summary distributions.

When a metric had a unimodal density, subject-level quartiles after robust standardization (median/MAD) along with uncertainty quantification using nonparametric bootstraps across subjects (2,000 resamples) were used to evaluate and determine high and low cognitive demand thresholds. The 25^th^ and 75^th^ percentiles were used as cut points and evaluated with the bootstrap 95% confidence intervals difference didn’t cross 0 and <2 and a separation d > 1.

Because 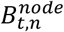 is a probabilistic posterior describing node-level at time *t*, a threshold was not applied. Rather, where further description of the most probable spatially distinct nodes that are responding at each timepoint is desired (e.g., counts or spatial overlays), an adaptive FDR correction was tested. Given the high-dimensional setting with a diffuse posterior mass it was critical to balance specificity while ensuring a sensitivity retention. Given that FDR alphas can be set for specific use cases, a range of alphas for FDR were examined to confirm that substantive results are stable over α *ϵ* [0·10, 0·30].

#### 2.4.2 Scaling Factors

Where processing mode and diagnostic metrics are already on a comparable scale in and of themselves, they operate on different scales resulting in variability in derivative metrics (i.e., demand, resource level, and bottleneck). Simulations were used to empirically derive scaling factors that stabilize these metrics in a common space for both subject-level and cross-study analysis/comparisons. Specifically, for each metric, the median and MAD-based Gaussian consistency factors were extracted across all subjects that are used for rescaling values, where GCC = Gaussian consistency constant (1.4826) then: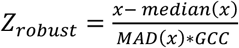 . This standardization procedure ensures comparability of emergent PCSM metric across different datasets while preserving their relative within-subject dynamics.

## 3 Results: Simulation Study –Ground Truth Recovery to Validate PCSM Alignment

### 3.1 Verifying Model Parameters and Priors

#### 3.1.1 GMM-HMM Model Parameters

The parsimonious model that met out-of-sample prediction criteria for generalizability was the 2 mixture and 3 latent states. The 2 mixture and 3 latent state model had the most substantive improvement in loglikelihood *Δ*_*LL*_= 3.47 with low standard error (0.013) demonstrating its utility for the GMM-HMM model (Table 2).

#### 3.1.2 Global Model for Individual Empirical Priors

Log-likelihoods were stable across folds, with all values (range: −114.595 to −113.513) falling within two standard deviations of the mean (−113.956 ± 1.278; SD = 0.639). Spatial similarity was also high: Dice coefficients indicated strong consistency between fold-specific and global models for both BOLD means (0.977 ± 0.040) and variances (0.934 ± 0.026). These results confirm that the GMM-HMM captures consistent spatial activation patterns across resampling folds, supporting model generalizability and parameter stability.

### 3.2 Parameter Recovery of Latent States and Transitions

Latent state recovery across all subjects ranged between 98.2%−99.9% for all states. This high recovery rate reflects identifiability and state-alignment accuracy under known generative conditions and is expected, given the well-established recovery properties of GMM-HMMs when the inference model matches the data-generating process; these values should not be interpreted as predictive performance on empirical data. Overall Accuracy was 99.1% with a precision of 99.4%, recall of 99.3% and a f1-score of 98.9% (Table 3, Figure 3). Recovery remained high across all noise levels and all transition probabilities, noting only minimal degradation for low transition probability simulations (average f1-score 0.999 for high and medium; 0.968 for low). Transition recovery also remained high with no drops by noise or transition probability to report. This indicates that model alignment and decoding are robust to both stochastic and deterministic variation in BOLD inputs.

**Figure 3.**
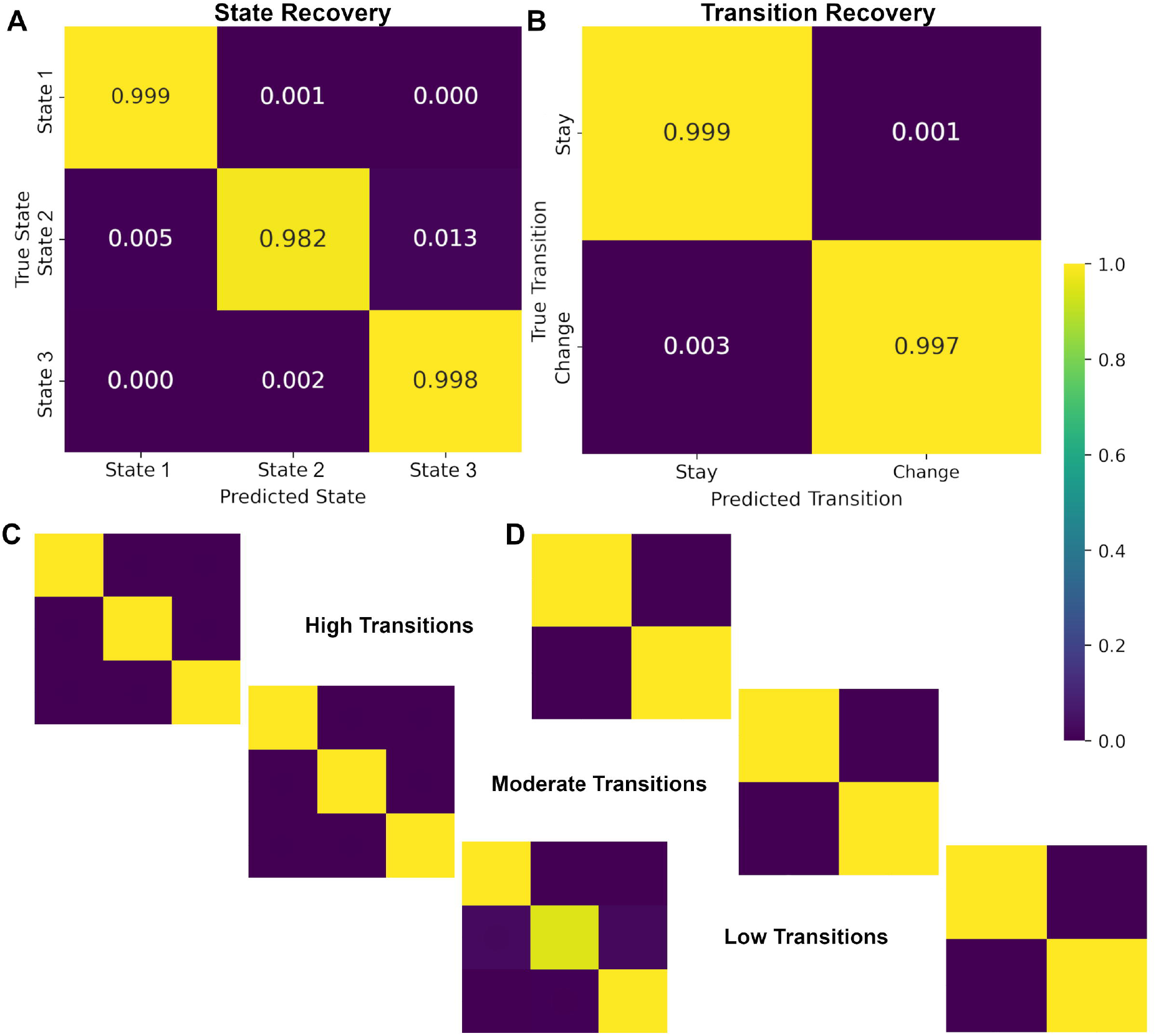

Together, these results confirm that alignment to the global GMM-HMM template yields stable, reproducible decoding of latent brain states suitable for downstream PCSM metrics. The high recovery accuracy demonstrates that PCSM can identify latent states with high certainty at each timepoint, even under noisy or rapidly switching conditions.

## 4 Metric Thresholding Procedures

### 4.1 Diagnostic Metrics

#### *ρ*_*t*_ Thresholding

Gaussian mixture modeling identified an upper threshold (τ= 0.991) with a strong separation (d= 27.914). Although two gaussian mixtures were identified, the lower distribution had < 20% of the density weight resulting in a threshold on the upper end of this metrics distribution. Accordingly, ρ_*t*_ ≥ 0.991 indicates strong temporal consistency whereas values under this threshold indicate weak temporal consistency indicating less reliable decoding and consideration for removal.

Sensitivity analyses across simulated transition probabilities confirmed expected trends: high transition probabilities yielded a greater proportion of weak temporal-consistency trials, whereas low transition probabilities yielded the highest proportion of strong-consistency trials (Table 4, Supplemental Table S1; Supplemental Figure S1).

#### *P*_*t*_ Thresholding

Gaussian mixture modeling identified a single dominant mode with a boundary-corrected KDE intersection (τ= 0.152) with a strong separation (d= 12.146). Thus, a *P*_*t*_ > 0.152 indicates strong BOLD response and cognitive engagement indicating more reliable decoding whereas values below this threshold indicate weak BOLD response and cognitive engagement that raise concerns for subsequent decoding.

Sensitivity analyses by transition probability revealed expected trends: High transition probabilities produced slightly more weak trial proportions (indicative of rapid, short-lived responses), whereas low transitions showed more sustained high-engagement trials (Table 5, Supplemental Table S1; Supplemental Figure S2).

#### 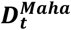 Thresholding

Gaussian mixture modeling identified a robust intersection threshold (τ=0.741) with a strong separation (d= 2.129). A 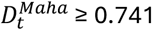 are flagged for further investigation with ρ_*t*_ and *P*_*t*_ because it indicates a trial with greater deviation from expected BOLD and therefore greater uncertainty in state decoding. Lower values indicate an expected range and more stable decoding.

Sensitivity analyses across transition probabilities (Table 6, Supplemental Table S1; Supplemental Figure S3) showed stable flagging rates (∼100 trials per condition), consistent with the expectation that deviations in BOLD intensity should not systematically vary with transition probability. These results support 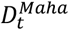 is a reliable cross-metric diagnostic for evaluating timepoints with potential decoding instability.

#### Diagnostics: Timepoint and Trial Flagging

Diagnostic analyses identified timepoints and trials with potential decoding instability, revealing expected patterns across simulations. Across the full sample, timepoints that met flagging criteria of 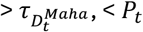, and < ρ_*t*_ averaged around 22.575±17.903. Of these flagged timepoint, only a fraction occurred more than once within a trial (0.002±0.024) with a range of 0-3 trials meeting removal criteria of ≥ 50% of flagged timepoints in a trial.

Sensitivity analyses by transition probabilities revealed high transitions had most of the flagged timepoints, but low transitions had the most flagged trials (Table 7). Because higher transition rates were expected to have higher higher single timepoint deviations that stabilize within trial and low transition rates were expected to have less single timepoint deviations but when they did occur, they would persist longer in a trial – this diagnostic measure behaves as expected. These results support that deviations from expected are a feature and indicate dynamic and short transitions. Importantly, flagged timepoints and trials represented a very small fraction of the total data, confirming that decoding stability across time and trials was well within acceptable bounds. Observed deviations reflect genuine neural transition dynamics rather than model error, supporting interpretability of subsequent PCSM measures.

### 4.2 Processing Mode Metrics

#### 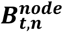 Adaptive FDR

Examining alphas *a* = *ϵ* [0·10, 0·30] revealed that *a* = 0·25 provided a balanced trade-off between sensitivity and sparsity (Table8). At α = 0·25, the realized false-discovery rate was conservative 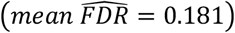, with an average of ∼ 11 nodes retained per timepoint. This corresponds to an expected ∼2 false nodes per timepoint. Increasing to *a* = 0·30 raised the average retained nodes to ∼18 with a lower probability threshold (*mean* τ = 0· &#x03C4;), and, although improving precision, decreasing *a* induced sparsity at the expense of sensitivity. Therefore *a* = 0·25 was adopted as a default for describing individual node responses per timepoint (Table 7).

Critically, this FDR-threshold map is only used for describing the spatial contributions of responding per timepoint. Serial-parallel determinations are not based on this FDR threshold; rather, inferences on serial or parallel processing are based on the full probability distribution of 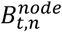 as the input for decoding cognitive processing states in 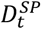.

#### 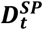 Thresholding

Gaussian mixture modeling identified two candidate intersections τ_*low*_ = 0·542 and τ_*high*_ = 0·612. The high intersection met separation criteria (d = 3.149) and was retained as a hard cutoff, where 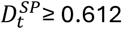 indicate serial processing. The low intersection, however, did not meet separation criteria (d = 0.937), suggesting an overlap between parallel and mixed processing. Applying boundary-corrected KDE refinement shifted the low threshold down to τ_*low*_ = 0·522 within the density valley, which had a strong separation (d = 3.724). This is a much clearer distinction between parallel and mixed states. As a result, 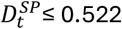 indicates strict parallel processing, while values between 0.522 and 0.612 represent mixed processing, and 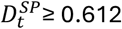 indicate serial processing.

Threshold quality analyses varying histogram resolution and kernel bandwidth produced highly stable estimates, with drift < 0.001 across all settings. Thus, the final thresholds were defined at 0.522 (low) and 0.612 (high), balancing mixture model structure with observed density distributions and features.

Sensitivity across transition probabilities was consistent with expected patterns. Specifically, a larger number of trials were serial due to completing cognitive functions tied to specific functions and the bottleneck. Transition specific mixed and parallel were as follows: a high transition probability had higher parallel and lower serial processing timepoints than those medium or low transition probabilities. Low transition probability simulations having the highest serial and lowest parallel processing timepoints as expected (Table 9, Supplemental Table S1; Supplemental Figure S4).

### 4.3 Demand and Resource Metrics

#### 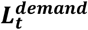 and 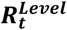 Thresholding

For 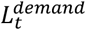, high and low thresholds (tau high = 0.675. 95%CI [0.674, 0.676]; tau low= −0.749, 95%CI [−0.752, −0.748]) met criteria for confidence as well as a strong separation (d = 17.409). Similarly, 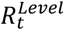 high and low thresholds (tau high = 0.595. 95%CI [0.594, 0.596]; tau low= −0.700, 95%CI [−0.704, −0.696]) met criteria for confidence as well as a strong separation (separation d = 7.381).

Sensitivity analyses examining transition probabilities confirmed theoretically consistent patterns: High transition probabilities were associated with more frequent high-demand timepoints and fewer high-resource timepoints, consistent with increased processing transitions. Conversely, low transition probabilities showed fewer high-demand timepoints but more high-resource intervals, reflecting sustained processing stability (Tables 10, 11, Supplemental Table S1; Supplemental Figures S5, S6)

### 4.4 Serial Bottleneck

#### *S*^*bottleneck*^ Thresholding

The serial bottleneck metric followed a trimodal distribution, thus a KDE approach was used to determine thresholds, the same as 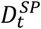. Two thresholds were identified along the upper tail >0 collectively representing about 17.1% of all values. The remaining 82.9% were < 0, representing the build of the distribution. The two thresholds represent long and extreme bottleneck (τ_*long*_ *≈* 0, τ_*extreme*_ *≈* 0·932) as they represented the 75^th^ and 90^th^ percentiles of the distribution in a tail. The long and extreme cut points had strong separation (long d= 8.635, extreme d= 2.691). Where the long distribution corresponds to 11.9% of the density, while the extreme only applied to 5.1% of the distribution representing the rarest and most persistent serial bottlenecks. Although τ_*extreme*_ captures extreme persistence, τ_*long*_ is recommended as the standard cutoff for general use due to its broader empirical representation and stability.

Sensitivity analyses across transition probabilities confirmed expectations: Low transition-probability simulations produced the greatest proportion of long and extreme bottlenecks, consistent with fewer but more sustained serial episodes; High transition probabilities, in contrast, produced more frequent but shorter serial periods (Table 12, Supplemental Table S1; Supplemental Figure S7).

### 4.5 Example Use of Metrics and Thresholds

Example outputs for high, medium, and low transition simulations can be found in supplemental materials (Supplemental Figures S8-S10). An example with OpenNeuro human fMRI data can be found in the GitHub Repository in notebooks.

## 5 Discussion

### 5.1 Conceptual Summary

The present work introduces Probabilistic Cognitive State Modeling (PCSM) that leverages posteriors from latent temporal and spatial modeling of human fMRI data to quantitate emergent cognitive properties during cognitive tasks. By providing trial-resolved timing data and aligning latent states and mixtures to redress GMM-HMM stochasticity, PCSM demonstrates a strong rate of recovery from learned dynamic patterns in human fMRI data – an essential foundation for robust emergent decoding.

From these posterior dynamics, PCSM derives interpretable emergent cognitive properties, including probabilistic node-level responses, serial–parallel processing, cognitive demand, and serial bottleneck persistence. Because these computational metrics lack external ground truth, their evaluation centers on identifiability, noise tolerance, stability under known generative conditions, and theoretical consistency; feasibility is demonstrated through stable estimation in simulations and successful application to empirical task data; and interpretability through formal metric definitions, predictable behavior in simulations, and coherent temporal structure in real data. Sensitivity is therefore defined in computational terms—reliable recovery and controlled degradation with noise—rather than via behavioral associations. The present work provides methodological validation and framework definition, establishing a necessary foundation for subsequent studies examining behavioral, developmental, clinical, or between-subject differences, which are important downstream applications rather than prerequisites for metric validity. Consistent with this aim, the metrics exhibit expected behavior and stable recovery across extensive simulation regimes.

Importantly, PCSM scales and normalizes these emergent metrics for robust and reliable cross-subject and cross-study comparisons. This enables PCSM to move beyond sample specific inferences to comparable value across samples, studies, and applications.

Together these tests bolster confidence in the value of PCSM as an approach to move away from static interpretations by affording tests of dynamic properties of cognition, cognitive processing, cognitive load, and serial bottlenecks that are well established as important features for understanding a variety of medical and mental health conditions.

### 5.2 Impact Summary

Methods such as PCSM provide a clear advantage by modeling individual-specific, dynamic and emergent properties of human cognition. It has been well established that cognitive processing and load demands are dynamic [4, 5] with individual-specific function and limits. These individual features are critical features of everyday functioning and psychiatric symptoms [13-18] that reflect variability rather than static deficits [19]. PCSM directly operationalizes this perspective by quantifying within-individual fluctuations in processing mode and demand – providing an approach to accomplish individual-specific investigation into cognition.

PCSM is a methodological advancement that links contemporary cognitive neuroscience with a probabilistic modeling framework for quantitating emergent properties of cognition. Where prior approaches have relied on static classifications, specialized paradigms, and low-dimensional data [30], PCSM leverages dynamic fMRI data to infer cognitive properties that can be applied across paradigms and task types.

This innovation builds direction on established and empirically supported contemporary models. Specifically that brain states follow a gaussian process [56], spatially distributed BOLD signals indexes distributed processing [32], and that cognitive transitions impose demands [9-11] that pull form cognitive resources with individual-specific limits [7, 8]. PCSM integrates this contemporary understanding into a unified probabilistic system that quantifies these dynamic and emergent properties over time.

By modeling the spatiotemporal and dynamic properties of fMRI data, PCSM fits a critical need in modern human neuroscience: to quantitate individual-specific cognitive variability, rather than average performance or static categories, as the meaningful signal. This approach supports a more nuanced and contemporary understanding of human cognition and its dysregulation.

### 5.3 Modeling Extensions

PCSM models time-resolved task-evoked activation patterns and does not model functional connectivity or causal interactions. However, the probabilistic outputs of PCSM (e.g., brain-state trajectories or node-level response probabilities) may be used as inputs to downstream analyses depending on the scientific question. For example, functional connectivity with B^node^ or using PCSM defined brain states as priors for Dynamic Causal Modeling [e.g., 65]. These extensions are conceptually straightforward but are outside the scope of the present work.

### 5.4 Conclusion

This project presents a robust approach to latent state modeling of fMRI data and decoding emergent cognitive properties critical for understanding adaptive and maladaptive cognition. PCSM demonstrates that the alignment of latent state organization and underlying BOLD mixture distributions show a strong consistency and recovery capturing most of the spatial and temporal structure. These results establish that PCSM produces posteriors representing dynamic features of the brain suitable for robust, comparable, and interpretable decoding of emergent cognitive properties.

Emergent decoding of processing, demand, resources, and serial bottleneck demonstrated stable relationships and theoretically expected patterns across transition probabilities and noise levels. Together, these findings demonstrate that PCSM effectively captures individual-specific fluctuations in control allocation and processing strategy, providing a bridge between computational models of cognition and human brain data.

By quantifying probabilistic transitions in processing and resource allocation, PCSM fills an important gap in cognitive modeling: linking moment-to-moment neural dynamics to cognitive processing and cognitive control theory. This framework positions PCSM as a scalable and generalizable method for decoding emergent cognitive phenomena in both healthy and clinical populations, advancing the broader goal of a probabilistic, dynamic study of human cognition.

## References

1. Logan, G.D., Parallel and serial processing. Stevens’ handbook of experimental psychology: Methodology in experimental psychology, 2002. 4: p. 271–300.

2. Townsend, J.T., Serial vs. parallel processing: Sometimes they look like Tweedledum and Tweedledee but they can (and should) be distinguished. Psychological science, 1990. 1(1): p. 46–54.

3. Townsend, J.T. and M. Fifić, Parallel versus serial processing and individual differences in high-speed search in human memory. Perception & Psychophysics, 2004. 66(6): p. 953–962.

4. Fischer, R. and F. Plessow, Efficient multitasking: Parallel versus serial processing of multiple tasks. Frontiers in psychology, 2015. 6: p. 1366.

5. Luria, R. and N. Meiran, Increased control demand results in serial processing: Evidence from dual-task performance. Psychological science, 2005. 16(10): p. 833–840.

6. Shenhav, A., et al., Toward a rational and mechanistic account of mental effort. Annual review of neuroscience, 2017. 40: p. 99–124.

7. Halford, G.S., W.H. Wilson, and S. Phillips, Processing capacity defined by relational complexity: Implications for comparative, developmental, and cognitive psychology. Behavioral and brain sciences, 1998. 21(6): p. 803–831.

8. Buschman, T.J., M. Siegel, J.E. Roy, and E.K. Miller, Neural substrates of cognitive capacity limitations. Proceedings of the National Academy of Sciences, 2011. 108(27): p. 11252–11255.

9. Band, G.P. and F.T. van Nes, Reconfiguration and the bottleneck: Does task switching affect the refractory period effect? European Journal of Cognitive Psychology, 2006. 18(4): p. 593–623.

10. Koch, I., E. Poljac, H. Müller, and A. Kiesel, Cognitive structure, flexibility, and plasticity in human multitasking—An integrative review of dual-task and task-switching research. Psychological bulletin, 2018. 144(6): p. 557.

11. Lien, M.-C., R. Schweickert, and R.W. Proctor, Task switching and response correspondence in the psychological refractory period paradigm. Journal of Experimental Psychology: Human Perception and Performance, 2003. 29(3): p. 692.

12. Meyer, D.E. and D.E. Kieras, A computational theory of executive cognitive processes and multiple-task performance: Part 2. Accounts of psychological refractory-period phenomena. Psychological review, 1997. 104(4): p. 749.

13. Meiran, N., G.M. Diamond, D. Toder, and B. Nemets, Cognitive rigidity in unipolar depression and obsessive compulsive disorder: Examination of task switching, Stroop, working memory updating and post-conflict adaptation. Psychiatry research, 2011. 185(1-2): p. 149–156.

14. Tillem, S., H. Weinstein, and A. Baskin-Sommers, Psychopathy is associated with an exaggerated attention bottleneck: EEG and behavioral evidence from a dual-task paradigm. Cogn Affect Behav Neurosci, 2021. 21(4): p. 881–893.

15. Winters, D.E., et al., Cognitive Control Difficulties Differentiate Callous-Unemotional Traits from Conduct Problems: A Pre-registered Double-blind Randomized Controlled Trial Analysis. 2025.

16. Winters, D.E., et al., Affective theory of mind impairments linked with callous-unemotional traits implicate cognitive control: A pre-registered double-blind randomized controlled trial with a dual-task paradigm. 2024.

17. Baskin-Sommers, A. and I.A. Brazil, The importance of an exaggerated attention bottleneck for understanding psychopathy. Trends in Cognitive Sciences, 2022. 26(4): p. 325–336.

18. Newman, J.P. and D.S. Kosson, Passive avoidance learning in psychopathic and nonpsychopathic offenders. Journal of abnormal psychology, 1986. 95(3): p. 252.

19. Allsopp, K., J. Read, R. Corcoran, and P. Kinderman, Heterogeneity in psychiatric diagnostic classification. Psychiatry research, 2019. 279: p. 15–22.

20. Miyake, A., et al., The unity and diversity of executive functions and their contributions to complex “frontal lobe” tasks: A latent variable analysis. Cognitive psychology, 2000. 41(1): p. 49–100.

21. Xu, T.L., K. De Barbaro, D.H. Abney, and R.F. Cox, Finding structure in time: Visualizing and analyzing behavioral time series. Frontiers in Psychology, 2020. 11: p. 1457.

22. Miller, J., R. Ulrich, and B. Rolke, On the optimality of serial and parallel processing in the psychological refractory period paradigm: Effects of the distribution of stimulus onset asynchronies. Cognitive psychology, 2009. 58(3): p. 273–310.

23. Dux, P.E., J. Ivanoff, C.L. Asplund, and R. Marois, Isolation of a central bottleneck of information processing with time-resolved FMRI. Neuron, 2006. 52(6): p. 1109–20.

24. Pashler, H., Overlapping mental operations in serial performance with preview. The Quarterly Journal of Experimental Psychology Section A, 1994. 47(1): p. 161–191.

25. Miller, E.K., S.L. Brincat, and J.E. Roy, Cognition is an emergent property. Current opinion in behavioral sciences, 2024. 57: p. 101388.

26. Gilman, J.M., J.M. Bjork, and T.E. Wilens, Brain signaling in psychiatric disorders: what can they tell us in the absence of behavioral differences? Journal of Behavioral and Brain Science, 2015. 5(08): p. 333.

27. Cohen, J.D., et al., Computational approaches to fMRI analysis. Nature neuroscience, 2017. 20(3): p. 304–313.

28. Sigman, M. and S. Dehaene, Brain mechanisms of serial and parallel processing during dual-task performance. Journal of Neuroscience, 2008. 28(30): p. 7585–7598.

29. Yildiz, A. and C. Beste, Parallel and serial processing in dual-tasking differentially involves mechanisms in the striatum and the lateral prefrontal cortex. Brain Structure and Function, 2015. 220: p. 3131–3142.

30. Li, K., et al., Distinguishing between parallel and serial processing in visual attention from neurobiological data. Royal Society Open Science, 2020. 7(1): p. 191553.

31. Taghia, J., et al., Uncovering hidden brain state dynamics that regulate performance and decision-making during cognition. Nature communications, 2018. 9(1): p. 2505.

32. Li, T., et al., Brain information processing capacity modeling. Scientific reports, 2022. 12(1): p. 2174.

33. Baker, A.P., et al., Fast transient networks in spontaneous human brain activity. elife, 2014. 3: p. e01867.

34. Vidaurre, D., S.M. Smith, and M.W. Woolrich, Brain network dynamics are hierarchically organized in time. Proceedings of the National Academy of Sciences, 2017. 114(48): p. 12827–12832.

35. Fincham, J.M., H.S. Lee, and J.R. Anderson, Spatiotemporal analysis of event-related fMRI to reveal cognitive states. 2020, Wiley Online Library.

36. Poldrack, R.A., et al., A phenome-wide examination of neural and cognitive function. Scientific data, 2016. 3(1): p. 1–12.

37. Glover, G.H., Deconvolution of impulse response in event-related BOLD fMRI1. Neuroimage, 1999. 9(4): p. 416–429.

38. Chen, G., et al., BOLD response is more than just magnitude: improving detection sensitivity through capturing hemodynamic profiles. Neuroimage, 2023. 277: p. 120224.

39. Lindquist, M.A., J.M. Loh, L.Y. Atlas, and T.D. Wager, Modeling the hemodynamic response function in fMRI: efficiency, bias and mis-modeling. Neuroimage, 2009. 45(1): p. S187–S198.

40. Taylor, A.J., J.H. Kim, and D. Ress, Characterization of the hemodynamic response function across the majority of human cerebral cortex. NeuroImage, 2018. 173: p. 322–331.

41. Taylor, A.J., J.H. Kim, and D. Ress, Temporal stability of the hemodynamic response function across the majority of human cerebral cortex. Human brain mapping, 2022. 43(16): p. 4924–4942.

42. Greene, A.S., et al., Why is everyone talking about brain state? Trends in Neurosciences, 2023. 46(7): p. 508–524.

43. Medaglia, J.D., et al., Brain state expression and transitions are related to complex executive cognition in normative neurodevelopment. NeuroImage, 2018. 166: p. 293–306.

44. Shirer, W.R., et al., Decoding subject-driven cognitive states with whole-brain connectivity patterns. Cerebral cortex, 2012. 22(1): p. 158–165.

45. Zhang, X., E.A. Maltbie, and S.D. Keilholz, Spatiotemporal trajectories in resting-state FMRI revealed by convolutional variational autoencoder. NeuroImage, 2021. 244: p. 118588.

46. Ezaki, T., Y. Himeno, T. Watanabe, and N. Masuda, Modelling state-transition dynamics in resting-state brain signals by the hidden Markov and Gaussian mixture models. European Journal of Neuroscience, 2021. 54(4): p. 5404–5416.

47. Baum, L.E., T. Petrie, G. Soules, and N. Weiss, A maximization technique occurring in the statistical analysis of probabilistic functions of Markov chains. The annals of mathematical statistics, 1970. 41(1): p. 164–171.

48. Viterbi, A., Error bounds for convolutional codes and an asymptotically optimum decoding algorithm. IEEE transactions on Information Theory, 2003. 13(2): p. 260–269.

49. hmmlearn, in Unsupervised learning and inference of Hidden Markov Models in python. 2024.

50. Bressler, D., N. Spotswood, and D. Whitney, Negative BOLD fMRI response in the visual cortex carries precise stimulus-specific information. PLoS One, 2007. 2(5): p. e410.

51. Wade, A.R., The negative BOLD signal unmasked. Neuron, 2002. 36(6): p. 993–995.

52. Wilson, R., A. Thomas, and S.D. Mayhew, Spatially congruent negative BOLD responses to different stimuli do not summate in visual cortex. NeuroImage, 2020. 218: p. 116891.

53. Scheeringa, R., et al., Neuronal dynamics underlying high-and low-frequency EEG oscillations contribute independently to the human BOLD signal. Neuron, 2011. 69(3): p. 572–583.

54. Everitt, B.S. and E.T. Bullmore, Mixture model mapping of brain activation in functional magnetic resonance images. Human brain mapping, 1999. 7(1): p. 1–14.

55. Hartvig, N.V. and J.L. Jensen, Spatial mixture modeling of fMRI data. Human brain mapping, 2000. 11(4): p. 233–248.

56. Shappell, H., B.S. Caffo, J.J. Pekar, and M.A. Lindquist, Improved state change estimation in dynamic functional connectivity using hidden semi-Markov models. NeuroImage, 2019. 191: p. 243–257.

57. Mitani, A., R. Sasaki, M. Oizumi, and T. Uka, A leaky-integrator model as a control mechanism underlying flexible decision making during task switching. PloS one, 2013. 8(3): p. e59670.

58. Drissi-Daoudi, L., A. Doerig, and M.H. Herzog, Feature integration within discrete time windows. Nature communications, 2019. 10(1): p. 4901.

59. Verbruggen, F., et al., A consensus guide to capturing the ability to inhibit actions and impulsive behaviors in the stop-signal task. elife, 2019. 8: p. e46323.

60. Schaefer, A., et al., Local-global parcellation of the human cerebral cortex from intrinsic functional connectivity MRI. Cerebral cortex, 2018. 28(9): p. 3095–3114.

61. Fabian, P., Scikit-learn: Machine learning in Python. Journal of machine learning research 12, 2011: p. 2825.

62. Davis, G., ATLAS: Mapping ATtention’s Location And Size to probe five modes of serial and parallel search. Attention, Perception, & Psychophysics, 2024. 86(6): p. 1938–1962.

63. Dice, L.R., Measures of the amount of ecologic association between species. Ecology, 1945. 26(3): p. 297–302.

64. Pedregosa, F., et al., Scikit-learn: Machine learning in Python. the Journal of machine Learning research, 2011. 12: p. 2825–2830.

65. Friston, K.J., et al., Dynamic causal modelling revisited. Neuroimage, 2019. 199: p. 730–744.

