## Supplemental for "Probabilistic Cognitive State Modeling (PCSM): Decoding Latent Spatiotemporal Dynamics to Reveal Serial-Parallel Processing, Cognitive Demand, and Serial Bottleneck in Task-Based fMRI"

Author: Drew E. Winters

**Supplemental Section 1: Parameterization**

Thresholds were derived from Gaussian mixture modeling (GMM) and kernel density estimation (KDE) applied to simulation-derived distributions for each metric. Intersections and valley minima were used to determine lower and upper boundaries, with separation quantified by Cohen's d.

Supplemental Table S1. Thresholds for metrics and separation

| Metric | $\tau_{low}$ | $\tau_{high}$ | Separation (d) | Notes |
|---|---|---|---|---|
| | Diagnostics | | | |
| $D_t^{Maha}$ | -- | 0.7409035409035409 | 2.128 | $> \tau_{high} = investigate$ |
| $P_t$ | 0.1524348304585251 | -- | 12.746 | $< \tau_{low} = flag$ |
| $\rho_t$ | -- | 0.8604972640007917 | 27.914 | $< \tau_{high} = flag$ |
| | Serial-Parallel | | | |
| $D_t^{SP}$ | 0.52233887187500 | 0.612202672131343 | [3.724, 3.149] | $< \tau_{low} = parallel$<br>$> \tau_{high} = serial$ |
| | Demand and Resources | | | |
| $L_t^{demand}$ | -0.749974752766904 | 0.675287330352096 | 17.409 | $< \tau_{low} = recovery$<br>$> \tau_{high} = demanding$ |
| $R_t^{level}$ | -0.700797686780789 | 0.595688706474272 | 7.381 | $< \tau_{low} = scarce$<br>$> \tau_{high} = more\ available$ |
| | Serial Bottleneck | | | |
| $S^{bottleneck}$ | -- | 0.0000010042364280 | [8.634, 2.691] | $> \tau_{high} = long\ serial\ bottleneck$ |

**Supplemental Section 2: Threshold Distributions and Visuals**

The following density plots show the simulated distributions used to define data-driven thresholds (τ) for diagnostic and decoding metrics in PCSM. For each metric, KDE curves visualize separation between regimes (e.g., low vs. high reliability or parallel vs. serial processing). Vertical dashed lines mark selected thresholds; labels report empirical values derived from simulation-based GMM–KDE estimation. Minor flattening near distribution boundaries (e.g., $D^{Maha}$) reflects expected boundary bias for non-negative variables and does not affect threshold validity.

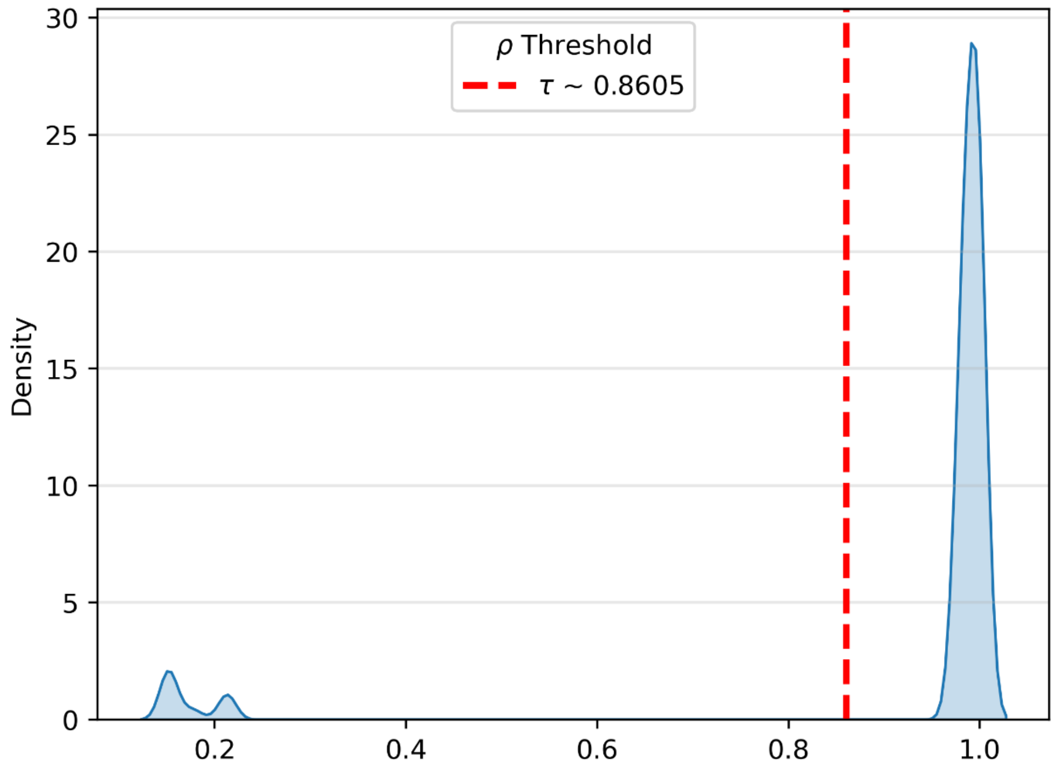


Figure S1. $\rho_t$ Threshold Density. Density distribution of temporal reliability ($\rho_t$) across simulations. The temporal reliability metric ($\rho_t \in [0,1]$) quantifies the stability of latent-state posterior probabilities between consecutive timepoints. Gaussian mixture modeling identified an upper threshold ($\tau \approx 0.991$) representing strong temporal consistency. Values below this threshold indicate reduced stability in latent-state decoding and are flagged for diagnostic consideration.

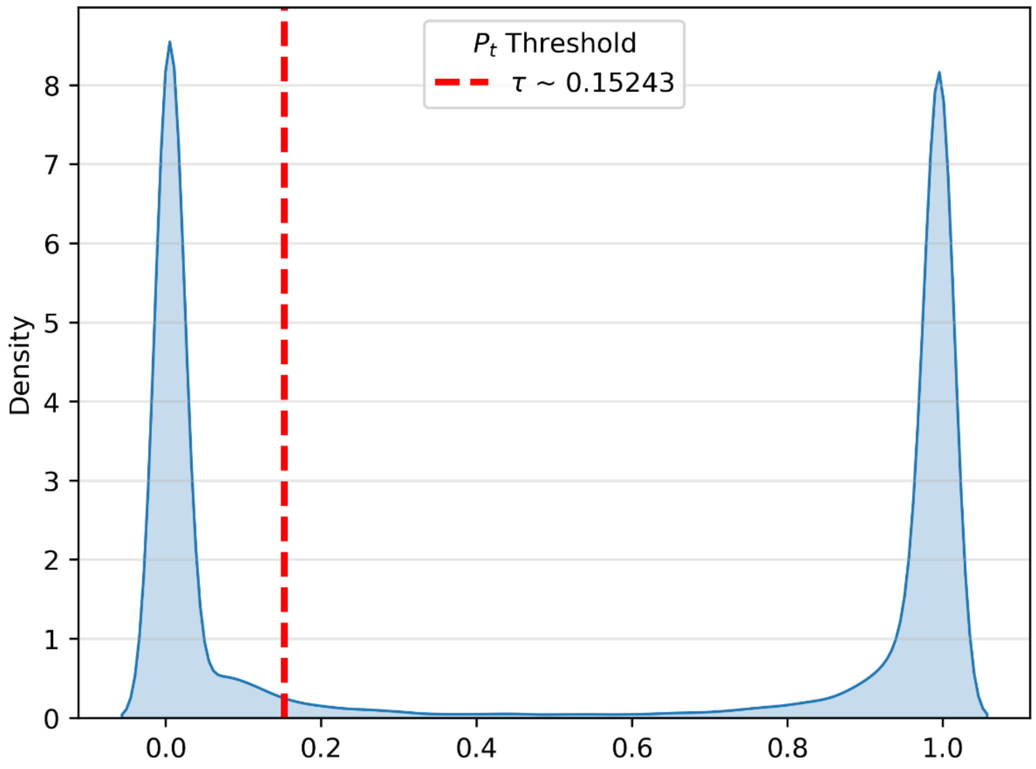


*Figure S2. $P_t$ Threshold Density. Density distribution of brain response probability ($P_t$) across simulations. The response probability metric ($P_t \in [0,1]$) quantifies the instantaneous probability that the brain is actively responding to task stimuli. Gaussian mixture modeling and boundary-corrected KDE refinement identified a threshold ($\tau \approx 0.152$) marking the boundary between weak and strong task-related engagement. Values above this threshold indicate strong BOLD response probability.*

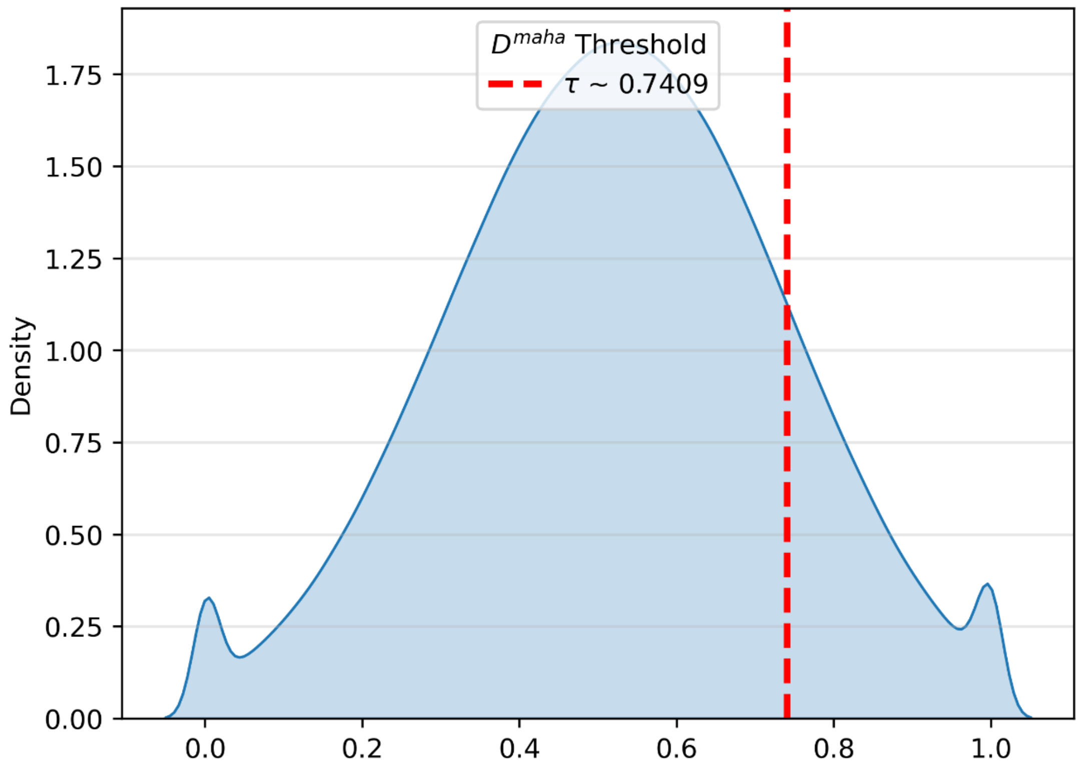


Figure S3. $D^{Maha}$ Threshold Density. Density distribution of Mahalanobis deviation ($D^{Maha}$) across simulations. The Mahalanobis distance ($D^{Maha} \geq 0$) quantifies the deviation between observed and model-expected BOLD activity. The empirical 85th percentile ($\tau \approx 0.741$) used as a diagnostic threshold for elevated model–data discrepancy. Minor flattening near zero reflects expected kernel boundary bias due to the non-negative support of $D^{Maha}$.

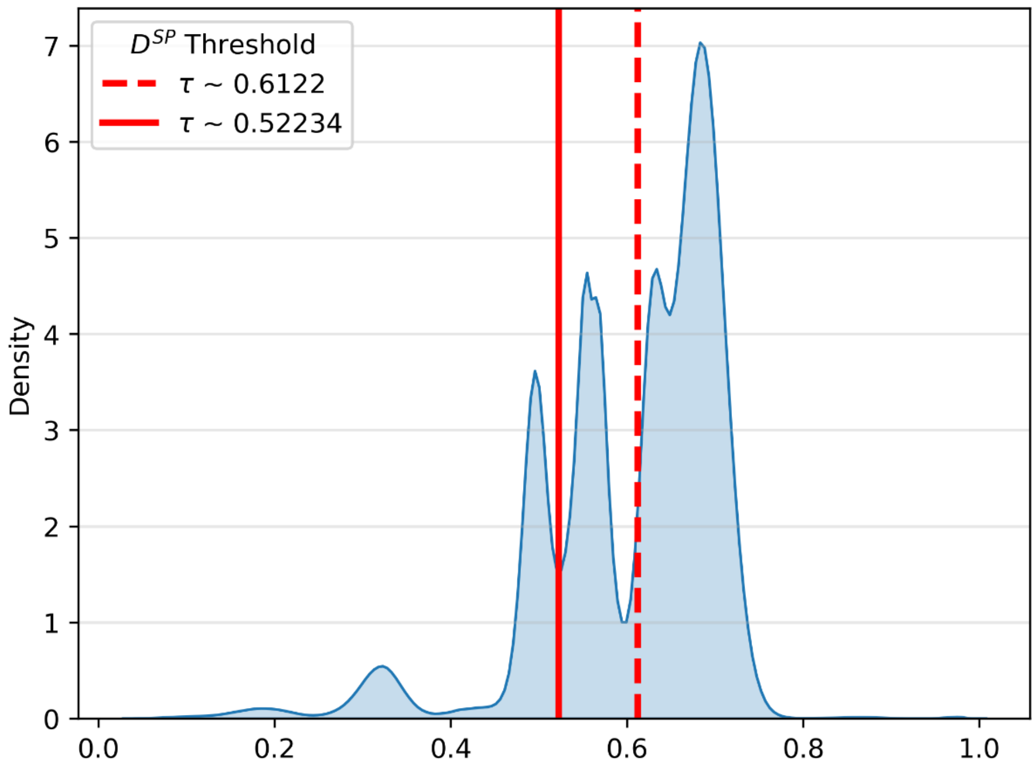


Figure S4. $D_t^{SP}$ Threshold Density. Density distribution of Serial–Parallel deviation index ($D_t^{SP}$) across simulations. $D_t^{SP}$ quantifies the average deviation in the number of responding nodes from half the total (2/N), distinguishing parallel (low) from serial (high) processing. KDE-based mixture modeling identified two data-driven thresholds (τ_low ≈ 0.522; τ_high ≈ 0.612) representing parallel and serial processing boundaries, respectively. Intermediate values indicate mixed processing modes.

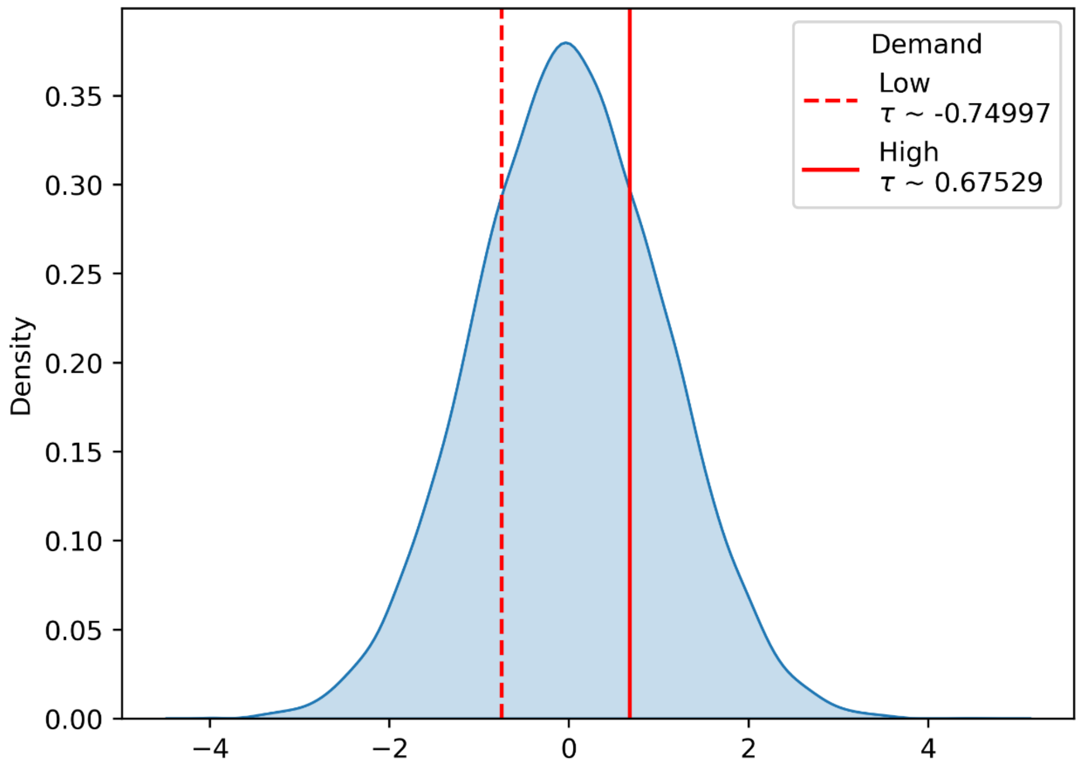


Figure S5. $L_t^{demand}$ Threshold Density. Density distribution of cognitive demand ($L_t^{demand}$) across simulations. $L_t^{demand}$ represents the signed moment-to-moment flow between processing strain and recovery. High and low thresholds (τ_high ≈ 0.675; τ_low ≈ −0.749) were defined by KDE intersection points, delineating periods of increased demand (negative values) versus recovery (positive values). The distribution shows near-symmetric variation around zero, reflecting balanced strain and recovery dynamics.

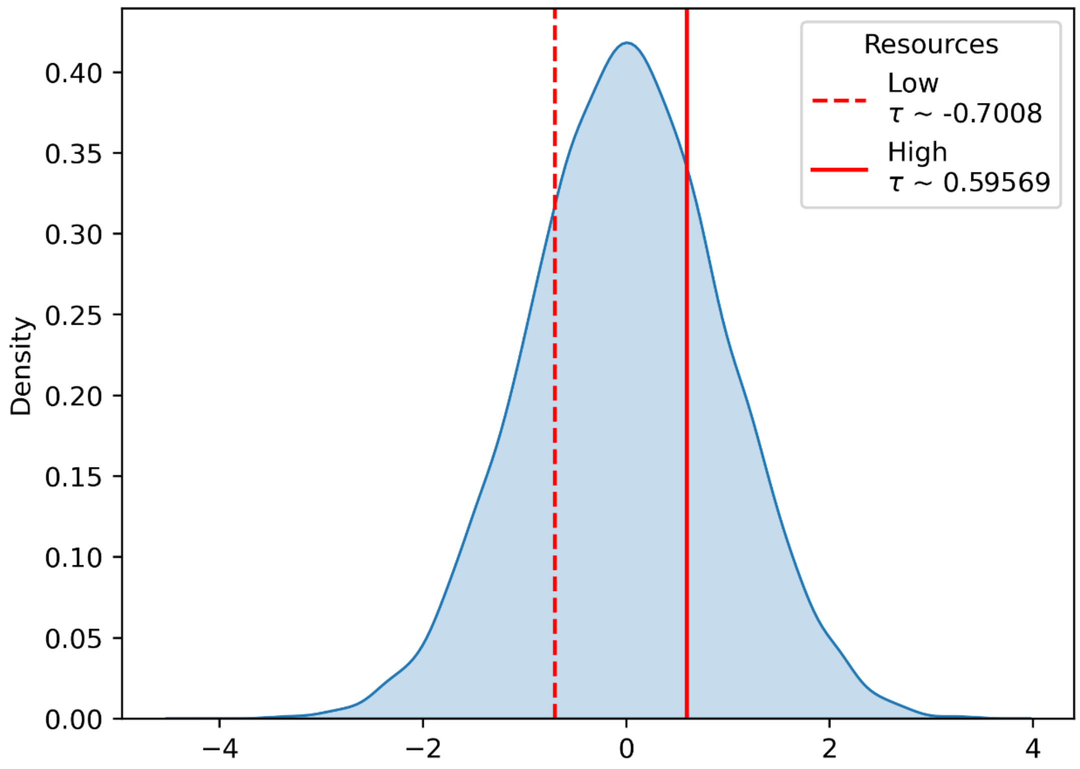


Figure S6. $R_t^{Level}$ Threshold Density. Density distribution of resource level ($R_t^{Level}$) across simulations. $R_t^{Level}$quantifies the cumulative availability of cognitive control resources as a leaky integration of demand over time. KDE-based thresholds (τ_high ≈ 0.595; τ_low ≈ −0.700) differentiate high-resource from low-resource states. The bimodal shape reflects alternating accumulation and depletion periods consistent with task-related control allocation.

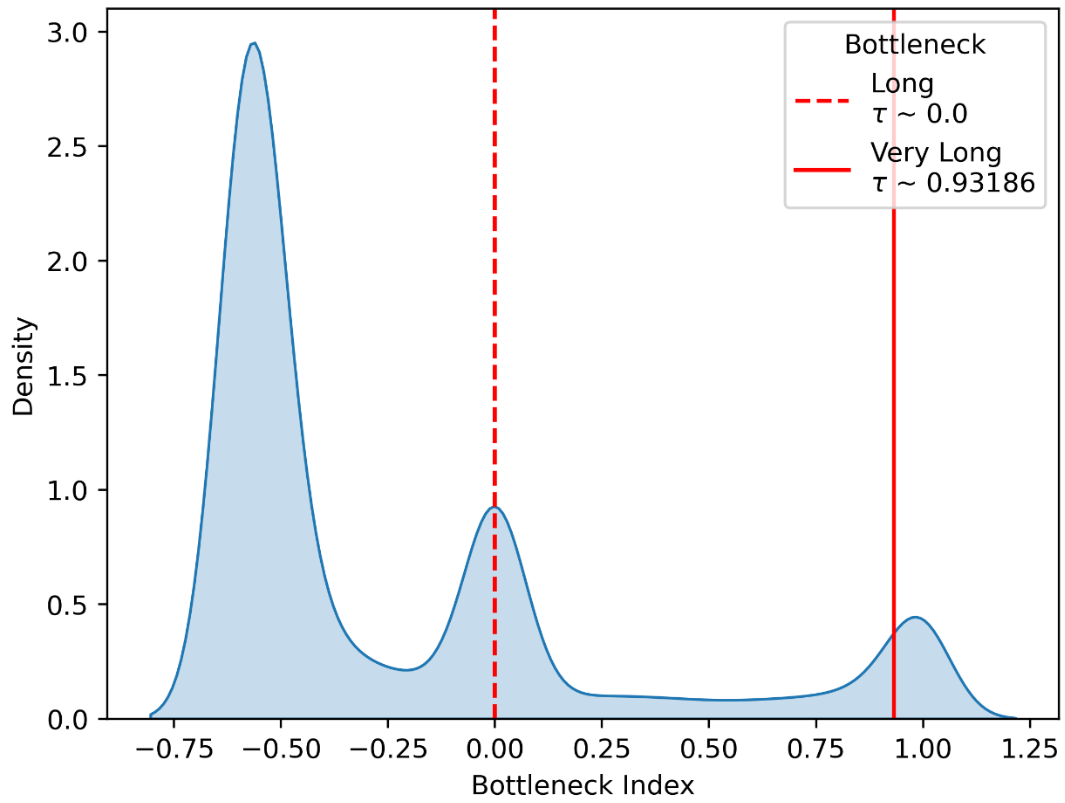


Figure S7. $S^{bottleneck}$ Threshold Density. Density distribution of serial bottleneck scalar ($S^{bottleneck}$) across simulations. $S^{bottleneck}$ measures the persistence of serial processing and effort required to disengage from serial episodes. KDE-based modeling of the trimodal distribution identified thresholds corresponding to long ($\tau \approx 0$) and extreme ($\tau \approx 0.932$) bottleneck persistence. Values below these cutoffs represent normal processing durations, while upper-tail extremes reflect high persistence and difficult disengagement.

Together, these distributions validate that simulation-based mixture and KDE methods yield stable, interpretable thresholds across diagnostic and cognitive decoding metrics

**Supplemental Section 3: Example Diagnostic and Transition Outputs**

The following panels illustrate how diagnostic metrics ($\rho_t$, $P_t$, and $D_t^{maha}$) and decoding metrics ($B_t^{Node}$, ${D_t}^{SP}$, ${L_t}^{demand}$, ${R_t}^{level}$, $S^{bottleneck}$) interact to evaluate model stability across simulated transition probabilities. High-, medium-, and low-transition regimes demonstrate consistent recovery of latent-state dynamics and expected trends in decoding reliability and processing mode classification.

### High Transitions

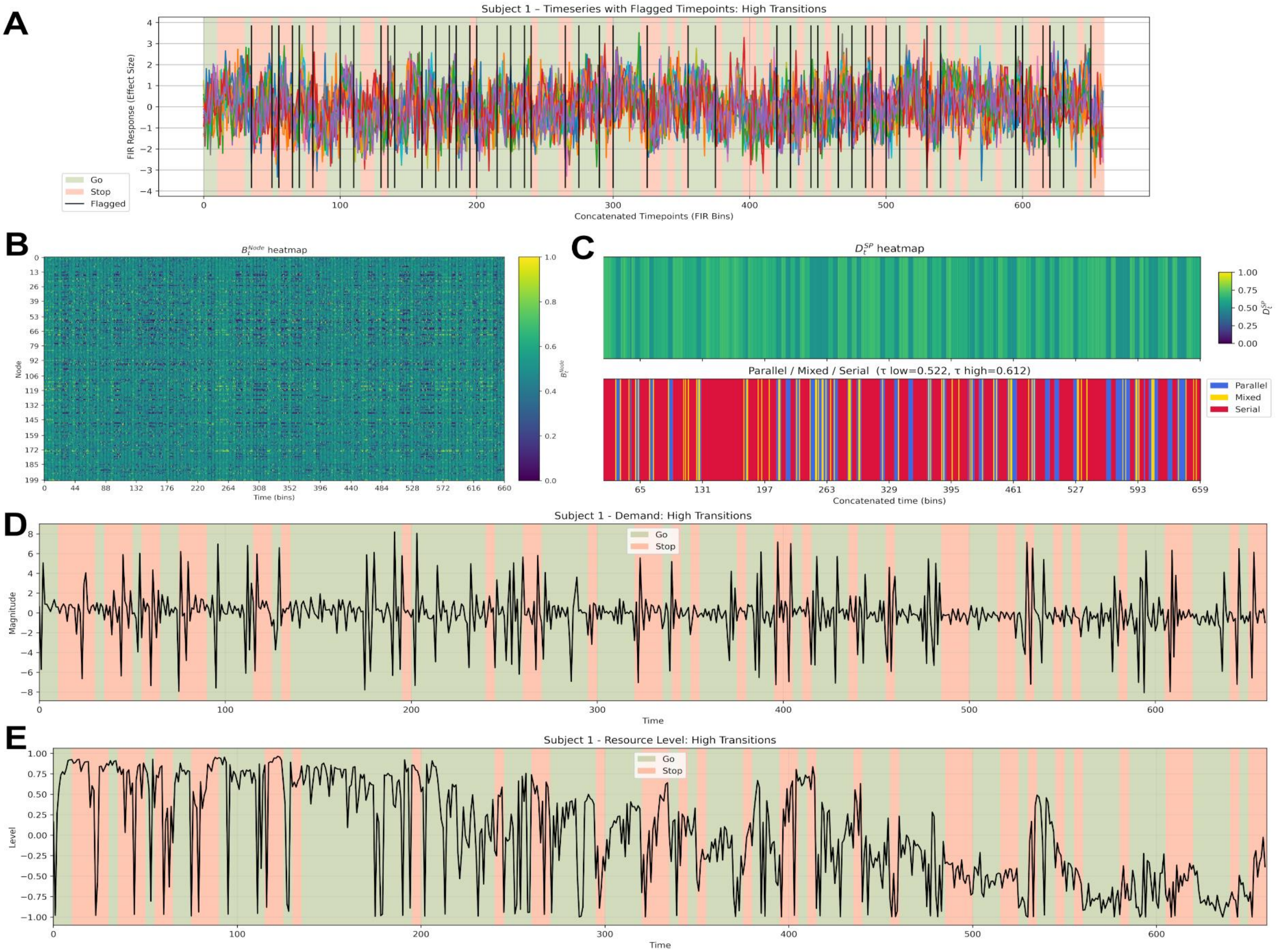


**Figure S8. Example outputs under the high-transition regime. A:** Simulated trajectories for diagnostic metrics ($\rho_t$, $P_t$, $D_t{}^{Maha}$) where lines indicate flagged timepoints – timepoints were not >50% within trial, **B**: node response ($B_t^{Node}$), **C**: serial–parallel processing ($D_t{}^{SP}$), **D**: cognitive demand ($L_t{}^{demand}$), and **E**: resource level ($R_t$^level). High-transition simulations show frequent latent-state switching, producing rapid alternations between serial and parallel processing, strong fluctuations in cognitive demand, and reciprocal resource-level dynamics.

**Medum Transitions**

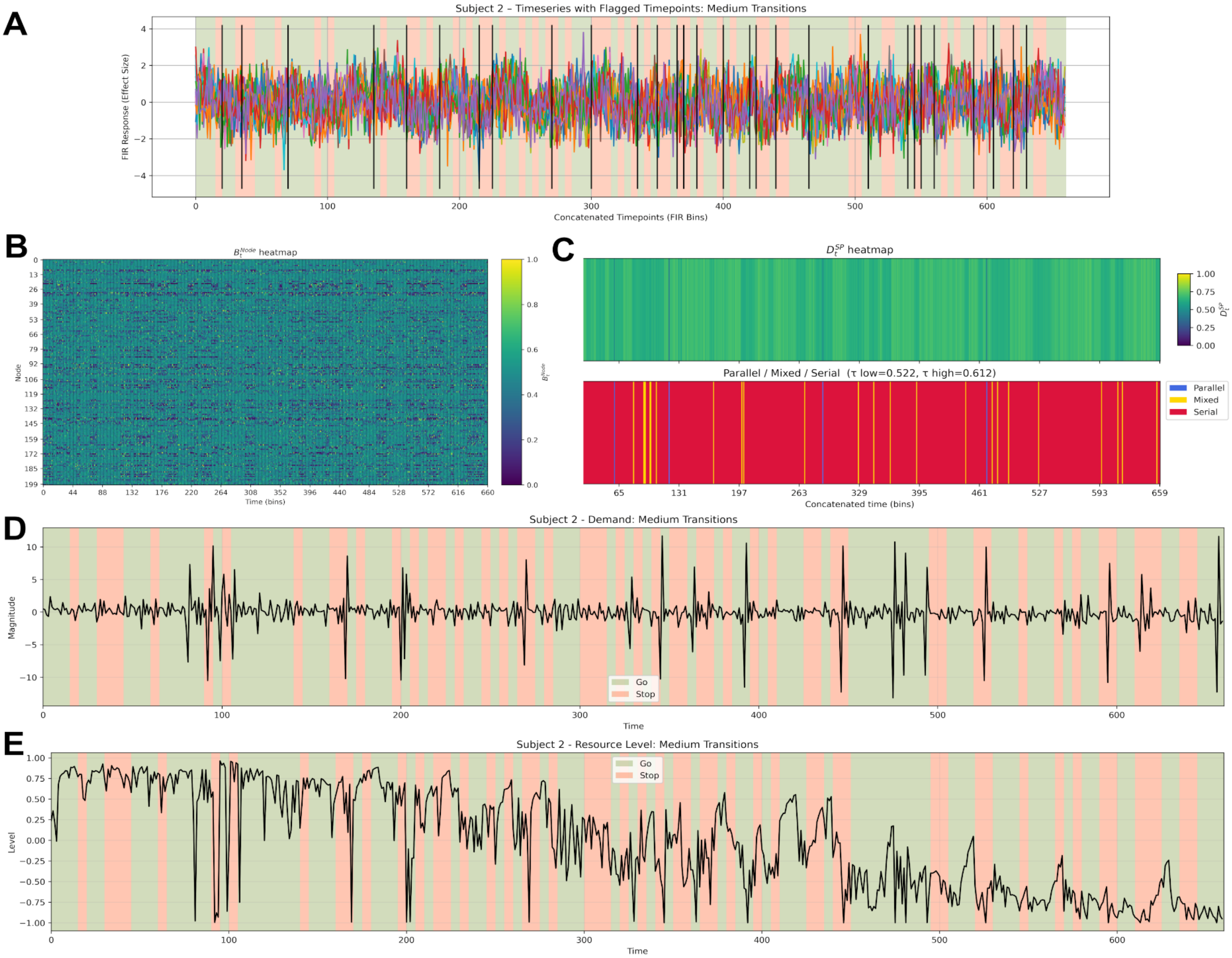


**Figure S9. Example outputs under the medium-transition regime.** Medium-transition simulations display moderate switching frequency with alternating but stable cognitive states. Demand and resource signals remain inversely coupled, and $D_t{}^{SP}$ shows balanced serial and parallel contributions over time.

### Low Transitions

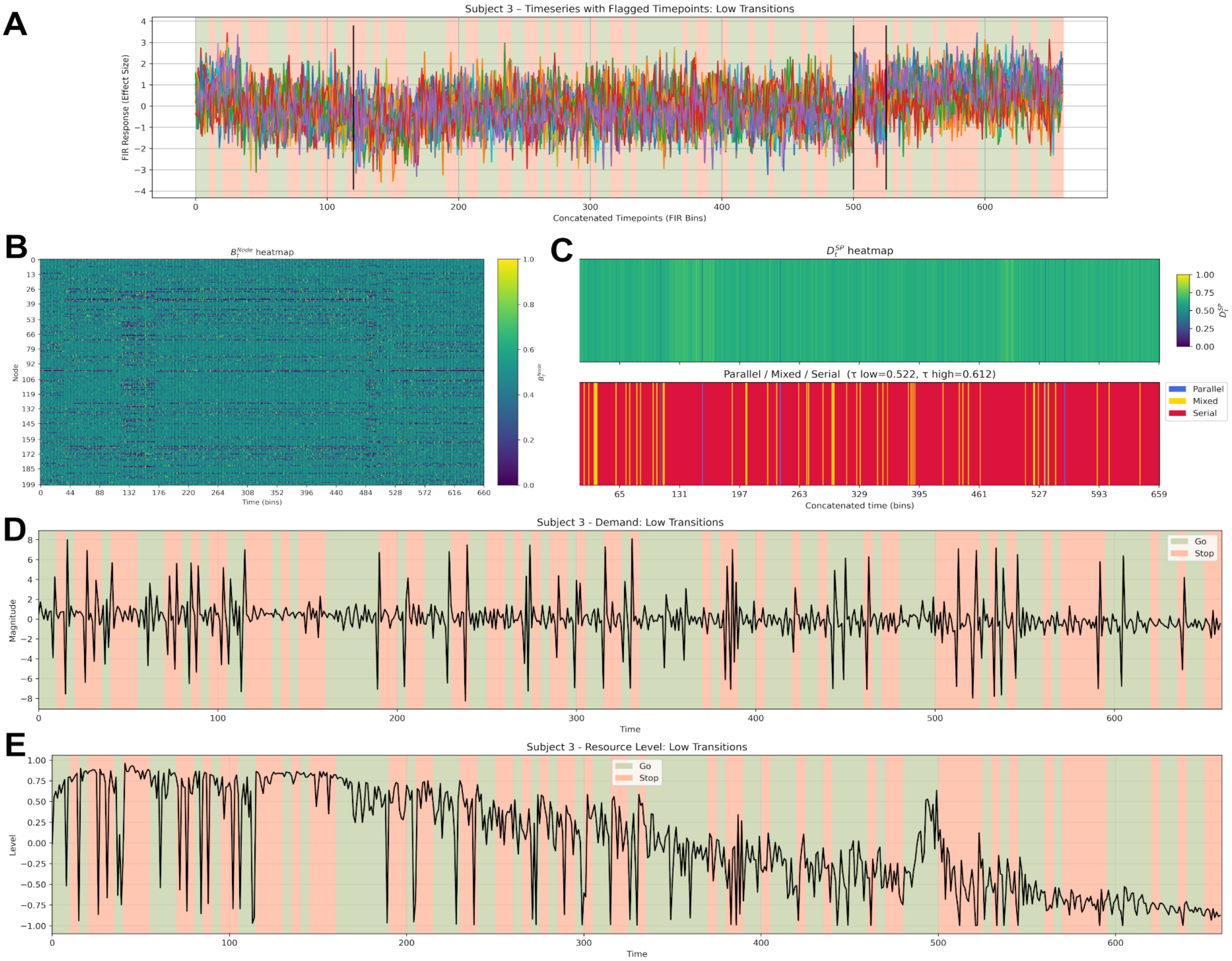


**Figure S10. Example outputs under the low-transition regime.** Low-transition simulations produce extended dwell periods with minimal state switching. Cognitive demand and resource level vary smoothly and inversely, and ${D_t}^{SP}$ remains near the parallel-mode boundary, consistent with sustained processing stability.
